# Exploring the Extreme Acid Tolerance of a Dynamic Protein Nanocage

**DOI:** 10.1101/2022.07.12.499790

**Authors:** Jesse A. Jones, Michael P. Andreas, Tobias W. Giessen

## Abstract

Encapsulins are protein nanocages capable of efficient self-assembly and cargo enzyme encapsulation. They are found in a wide variety of bacteria and archaea, including many extremophiles, and are involved in iron and sulfur homeostasis, oxidative stress resistance, and secondary metabolite production. Resistance against physicochemical extremes like high temperature and low pH is a key adaptation of many extremophiles and also represents a highly desirable feature for many biotechnological applications. However, no systematic characterization of acid stable encapsulins has been carried out, while the influence of pH on encapsulin shells has so far not been thoroughly explored. Here, we report on a newly identified encapsulin nanocage (AaEnc) from the acid-tolerant bacterium *Acidipropionibacterium acidipropionici.* Using transmission electron microscopy, dynamic light scattering, and proteolytic assays, we demonstrate its extreme acid tolerance and resilience against proteases. We structurally characterize the novel nanocage using cryo-electron microscopy, revealing a dynamic five-fold pore that displays distinct “closed” and “open” states at neutral pH, but only a singular “closed” state under strongly acidic conditions. Further, the “open” state exhibits the largest pore in an encapsulin shell reported to date. Non-native protein encapsulation capabilities are demonstrated, and the influence of external pH on internalized cargo is explored. AaEnc is the first characterized highly acid stable encapsulin with a unique pH-dependent dynamic pore and its molecular characterization provides novel mechanistic details underlying the pH stability of large dynamic protein complexes.

## Introduction

Protein-based compartments are used by many prokaryotes to regulate and optimize their metabolism in space and time.^1^ One of the most widespread families of microbial protein compartments are encapsulins. Structurally, encapsulins self-assemble from a single type of HK97 phage-like shell protein to form icosahedral nanocages.^2^ Encapsulins can exhibit different triangulation numbers, T=1 (60mer, ca. 24 nm), T=3 (180mer, ca. 32 nm), and T=4 (240mer, ca. 42 nm), with pores of varying sizes located at the 5-, 3-, and 2-fold symmetry axes.^3,4^ They are classified into four distinct families, with Family 1 being the first discovered and most studied,^4^ and derive their name from their ability to encapsulate specific co-regulated cargo enzymes. In Family 1, encapsulation is mediated by conserved peptide sequences found at the C-terminus of all cargo proteins called targeting peptides (TPs).^5^ This feature allows encapsulins to perform a variety of biological functions, such as to act as sequestration chambers for dye-decolorizing peroxidases (DyPs), involved in combating oxidative stress;^5,6^ to serve as iron mineralization and storage compartments;^7–10^ as well as to sequester desulfurase enzymes, likely involved in sulfur metabolism.^11^

Due to their favorable properties, encapsulins have gained much attention as bioengineering tools.^3,12,13^ As such, engineered encapsulins have been used in bacteria, yeast, and human cells for various applications, including as metabolic nanoreactors,^14^ cellular imaging systems,^13^ and drug delivery platforms.^15,16^ A recent increase in the number of studies on encapsulin systems highlights the expanding scope of the field.^2,4,10,11,17^ The encapsulin shell in particular has received substantial attention in recent years.^17^ Efforts aimed at increasing shell stability,^15^ controlling shell assembly,^18^ and modulating pore size and dynamics have recently been reported.^19,20^ Encapsulin shells efficiently self-assemble under many conditions and display marked resistance against chemical or temperature denaturation, pH, and non-specific proteases. For example, the melting temperature of the T=4 shell from *Quasibacillus thermotolerans* was reported to be nearly 87°C,^10^ while the encapsulin from *Brevibacterium linens* was shown to be stable across a broad range of pH values (pH 5 to 11).^21^ However, a number of biotechnological and industrial processes – including lignocellulose hydrolysis for biofuel production,^22^ breakdown of complex sugars for monosaccharide production,^22^ and bioleaching to prevent metal contamination and enable bioremediation in the mining industry^23,24^ – would benefit from modular protein cages stable at acidic conditions below pH 5. Even though evolutionary adaptations of thermostable proteins have been well characterized and include oligomerization, large hydrophobic cores, and disulfide bond formation, adaptations that lead to acid-stable proteins are poorly understood.^25,26^

Here, we carry out the first bioinformatic search for acid-stable encapsulin nanocages and subsequently characterize the structure and stability of a Family 1 encapsulin shell from *Acidipropionibacterium acidipropionici* (AaEnc) – one of the top hits identified in our *in silico* analysis. Using a combination of techniques, including cryo-electron microscopy (cryo-EM), we characterize the structure and acid stability of the AaEnc nanocage across a wide range of pH values. Our results highlight the acid stability of AaEnc and its resilience towards protease digestion under different pH conditions. Cryo-EM analysis reveals a pH-dependent dynamic 5-fold pore with defined “closed” and “open” states, with the latter representing the largest pore in an encapsulin shell reported to date. Further analyses confirm the non-native cargo loading capabilities of AaEnc and reveal the effects of external pH on internalized cargo.

## Results and Discussion

### Bioinformatic search for acid-tolerant encapsulin shells

Only Family 1 encapsulins have so far been used as engineering platforms as they are the most studied and well-understood of the different types of encapsulins – especially with respect to non-native cargo loading. Therefore, with future engineering applications in mind, we chose to focus our bioinformatic search for acid-tolerant shells on Family 1.^3,4,27,28^ Further arguments for focusing on Family 1 are that the widespread DyP encapsulins found in Family 1 are known to optimally function under acidic conditions^29^ while a variety of Family 1 systems encoded by acid-tolerant and acidophilic bacteria have already been previously identified.^4^

As acid-tolerant proteins generally possess a low calculated isoelectric point (pI) caused by a large number of surface-exposed negatively charged residues,^22,25,26,30^ it was hypothesized that encapsulins with a low pI may exhibit increased acid tolerance. Therefore, the large set of previously identified Family 1 encapsulins with molecular weights between 26 and 35 kDa – the size range of non-cargo fused encapsulins – were ranked by pI (**Figure 1a, Figure 1b,** and **Supplementary Data 1**). The lowest observed pls ranged from 4.14 to 4.49 with one protein found in the pI bin centered at a pI of 4.2 and 28 proteins found in the pI bin centered at a pI of 4.4. Of these encapsulins, 18 are encoded by halophiles, four by acidophiles or acid-tolerant bacteria, and the rest by soil bacteria or putative pathogens (**Table S1**).

**Figure 1.**
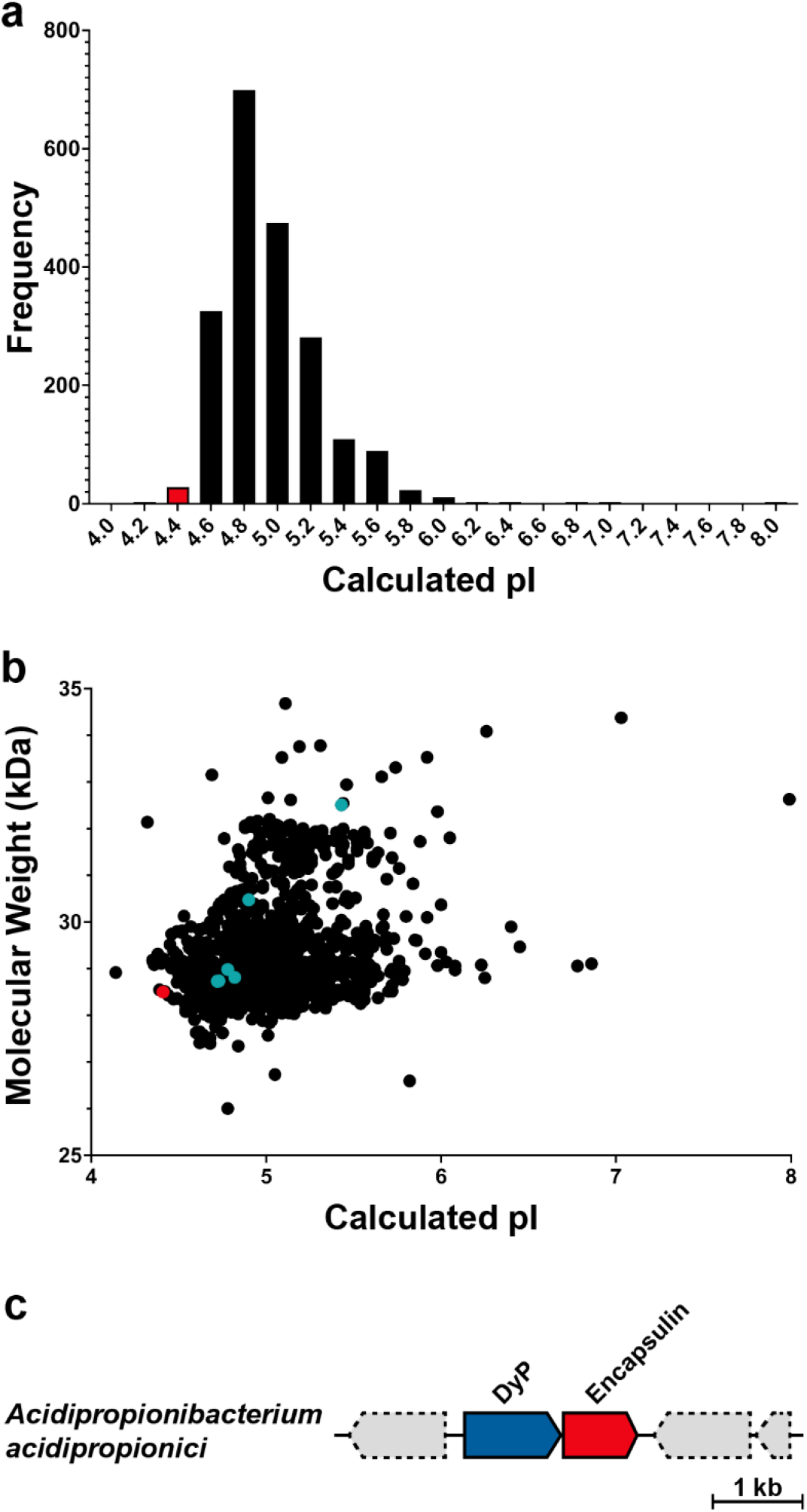
Bioinformatic analysis of Family 1 encapsulins. **a**) Histogram of encapsulins based on calculated isoelectric point (pI), binned at 0.2 pH units. The pI bin of interest centered at pI 4.4 and containing the *A. acidipropionici* encapsulin (AaEnc) is highlighted in red. **b**) Scatter plot of encapsulins based on calculated pI and molecular weight (MW). AaEnc (pI: 4.41, MW: 28.5 kDa) is shown in red. Previously well-characterized encapsulins from *Mycobacterium smegmatis, Brevibacterium linens, Mycolicibacterium hassiacum, Haliangium ochraceum, Thermotoga maritima,* and *Myxococcus xanthus* are shown in cyan. **c**) The *A. acidipropionici* Family 1 encapsulin operon containing a dye-decolorizing peroxidase (DyP) cargo enzyme (blue) and encapsulin shell (AaEnc, red). Functionally unrelated genes are shown in gray with dashed outlines. Scale bar: 1 kilobase (kb).

We chose to focus on an encapsulin encoded by one of the acid-tolerant species, namely, the DyP system of the industrially relevant *A. acidipropionici* (ATCC 4875) (AaEnc; **Figure 1c, Table S1,** and **Figure S1**). *A. acidipropionici* is a Gram-positive actinobacterium able to tolerate acidic conditions as low as pH 4.4.^31^ It is used in agricultural applications and studied for its biotechnological and industrial potential due to its production of propionic acid as a primary fermentation product, with acetic acid and carbon dioxide as secondary products.^31–33^

The pI and sequence composition of AaEnc was compared to previously characterized encapsulin shells, including those from *Mycobacterium smegmatis, Brevibacterium linens, Mycolicibacterium hassiacum, Haliangium ochraceum,^34^ Thermotoga maritima,* and *Myxococcus xanthus* (**Figure 1**, **Figure S1, Table 1,** and **Table S1**).^2,7,21,35,36^ AaEnc was found to have a lower pI than all of the so far characterized Family 1 encapsulins, and to display the largest ratio of acidic to basic residues (**Table 1** and **Table S1**). This makes AaEnc a promising test system for exploring the acid stability of encapsulin shells.

**Table 1.** Comparison of the pI and charge of AaEnc and other characterized encapsulins.

| Encapsulin | pI<sup>1</sup> | # Acidic residues | # Basic residues | Acid/base ratio | Charge at pH 7.0<sup>2</sup> |
| --- | --- | --- | --- | --- | --- |
| *A. acidipropionici* | 4.41 | 41 | 25 | 1.64 | -23.47 |
| *M. smegmatis* | 4.72 | 41 | 31 | 1.32 | -16.44 |
| *B. linens* | 4.73 | 40 | 30 | 1.33 | -18.77 |
| *M. hassiacum* | 4.78 | 41 | 32 | 1.28 | -15.50 |
| *H. ochraceum* | 4.82 | 40 | 32 | 1.25 | -13.47 |
| *T. maritima* | 4.90 | 50 | 39 | 1.28 | -13.05 |
| *M. xanthus* | 5.45 | 37 | 36 | 1.03 | -7.50 |

<sup>1</sup> Theoretical isoelectric points calculated with Expasy Compute pI/MW tool (https://web.expasy.org/compute_pi/).
<sup>2</sup> Charges at pH 7.0 calculated via Geneious Prime 2020.2.4 (https://www.geneious.com).

### Biophysical analysis of the AaEnc protein nanocage

To characterize the AaEnc nanocage, it was first heterologously expressed in *Escherichia coli* and then purified via a combination of polyethylene glycol (PEG) precipitation, anion exchange chromatography (IEC), and size exclusion chromatography (SEC), with the latter serving as an initial verification that the nanocage was assembled at pH 7.5 (**Figure 2a**). Aliquots of the purified sample were then exchanged into various buffers across a wide range of pH values (pH 1.5 to 10.0) while holding the salt concentration constant at a physiological value of 150 mM NaCl. After incubation for 6 h, samples were imaged via negative stain transmission electron microscopy (TEM) to assess the effect of pH on protein aggregation and the assembly state of the AaEnc nanocage. Some assembled AaEnc shells could be observed at pH values as low as 1.9 and as high as 7.5 (**Figure S2**). However, the pH range within which AaEnc was close to fully assembled with only minor aggregation occurring was between pH 2.25 and 7.5. Therefore, subsequent biophysical analyses were carried out at four pH values spanning this pH range, namely at pH 2.25, 3.0, 5.0, and 7.5. Dynamic light scattering (DLS) analyses and negative stain TEM indicated that AaEnc maintained a similar size and appearance across all four tested conditions with Z-average diameters of 30.3 nm at pH 2.25, 30.2 nm at pH 3.0, 29.8 nm at pH 5.0, and 24.4 nm at pH 7.5 (**Figure 2b** and **Figure 2c**). The slight increase in average diameter at acidic pH values is likely due to limited aggregation under these conditions, however, as can be seen in TEM micrographs, individual shells at all tested pH values exhibited diameters of ca. 24 nm. Static light scattering (SLS) further indicated that AaEnc is relatively stable across all tested pH values, with aggregation temperatures (T_agg_) of 36.0°C at pH 2.25, 38.3°C at pH 3.0, 39.3°C at pH 5.0, and 62.4°C at pH 7.5 (**Figure 2d**). However, a clear trend can be observed with lower pH values leading to decreased T_agg_ values. We next explored the resistance of AaEnc against proteolytic degradation at various pH values. AaEnc proved to be relatively resilient to pepsin degradation at pH 3.0 and 37°C during a 3 h incubation period, whereas the control protein, bovine serum albumin (BSA), was completely degraded under the same conditions (**Figure 2e**). Similarly, AaEnc was relatively resistant to degradation by trypsin and chymotrypsin at pH 7.5 and 37°C over an 8 h period, with BSA being again substantially degraded under the same conditions (**Figure 2f**). Overall, these results highlight the substantial acid stability of AaEnc which is significantly higher than that of any other previously characterized encapsulin.

**Figure 2.**
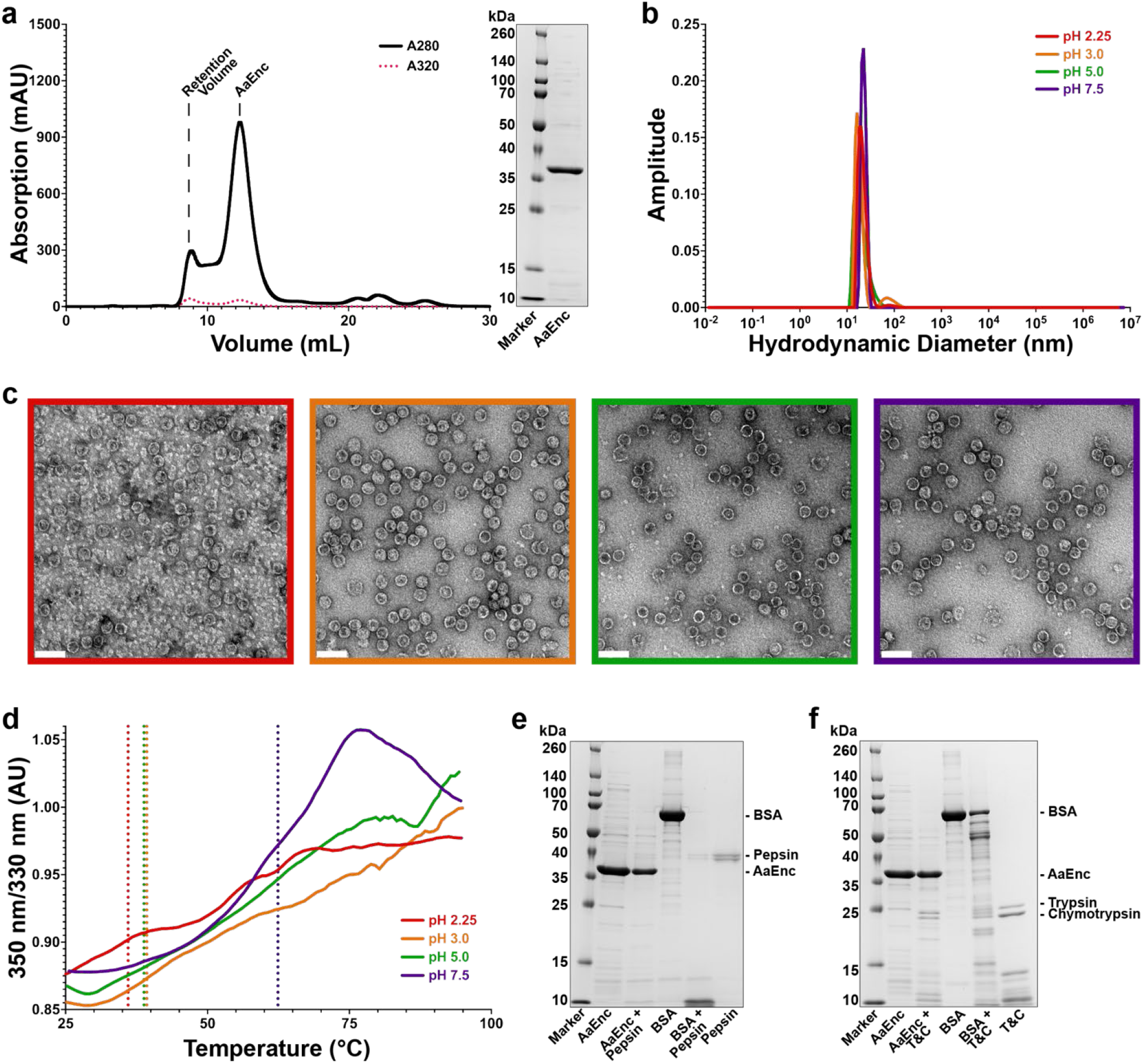
Biophysical analysis of the AaEnc nanocage. **A**) Size exclusion chromatography (SEC) of the AaEnc nanocage depicting elution at 12 mL suggestive of an assembled T=1 encapsulin (left), along with SDS-PAGE analysis of purified AaEnc (right). **B**) Dynamic light scattering (DLS) analysis of AaEnc at pH 7.5 (purple), pH 5.0 (green), pH 3.0 (orange), and pH 2.25 (red). **C**) Transmission electron microscopy (TEM) analysis of AaEnc after 6 h incubation at different pH values: pH 2.25 (red, far left), pH 3.0 (orange, middle left), pH 5.0 (green, middle right), and pH 7.5 (purple, far right). Scale bars: 50 nm. **D**) Representative thermal unfolding curves for AaEnc at different pH values: pH 2.25 (red), pH 3.0 (orange), pH 5.0 (green), and pH 7.5 (purple) with corresponding aggregation temperatures (T_agg_, vertical dashed lines, respective colors). **e**) Protease stability analysis of AaEnc exposed to pepsin at pH 3.0 for 3 h with bovine serum albumin (BSA) as a control. **f**) Protease stability analysis of AaEnc exposed to trypsin and chymotrypsin (T&C) at pH 7.5 for 8 h with BSA as a control.

### Structural characterization of the AaEnc protein nanocage

To further characterize the influence of pH on AaEnc, single particle cryo-EM analysis was carried out, initially at a physiological pH of 7.5. Results revealed the existence of two discrete structural states distinguished by an either all “closed” or all “open” conformation of the 5-fold pores within the AaEnc shell (**Figure 3** and **Supplementary Video 1**). The “closed” and “open” states were determined to 2.90 (29,056 particles) and 3.32 Å (13,581 particles), respectively (**Figure S3**). About 68% of the used particles comprise the “closed” state while about 32% exhibit an “open” state. The shell for both states consists of 60 AaEnc protomers, forming a ca. 1.7 MDa, T=1 icosahedral protein cage with a diameter of 24 nm and an overall negatively charged exterior surface (**Figure 3a** and **Figure 3c**). Symmetric (icosahedral, I) and asymmetric (C1) refinements were carried out for both states to investigate if a given pore can be “closed” or “open” independently of the other pores in the same shell or if all pore states are generally correlated. Both I and C1 refinements yielded similar all “closed” and all “open” states (**Figure S3**), thus confirming that under the given experimental conditions, the pore dynamics of all pentameric facets within a shell appear to be strongly correlated. However, this does not necessarily exclude the possibility that, under certain environmental conditions or in the presence of cargo, a single shell can contain both “closed” and “open” 5-fold pores at the same time. This phenomenon – independent dynamic 5-fold pores – has indeed been observed for the Familiy 1 encapsulin from *H. ochraceum* (**Figure S4**).^37^ To further analyze 5-fold pore dynamics, 3D variability analysis with three components was carried out in cryoSPARC for both datasets, however, no states could be resolved that would indicate the simultaneous presence of both “closed” and “open” pores within the shell. The 5-fold pore diameters in the AaEnc nanocage are 5 Å for the “closed” and 20 Å for the “open” state (**Figure 3b, Figure 3d**, and **Figure S3**). Thus, the AaEnc “open” state represents the largest pore found in an encapsulin shell to date, substantially larger than the previously reported dynamic *H. ochraceum* pore which exhibited an “open” state diameter of 15 Å.^37^

**Figure 3.**
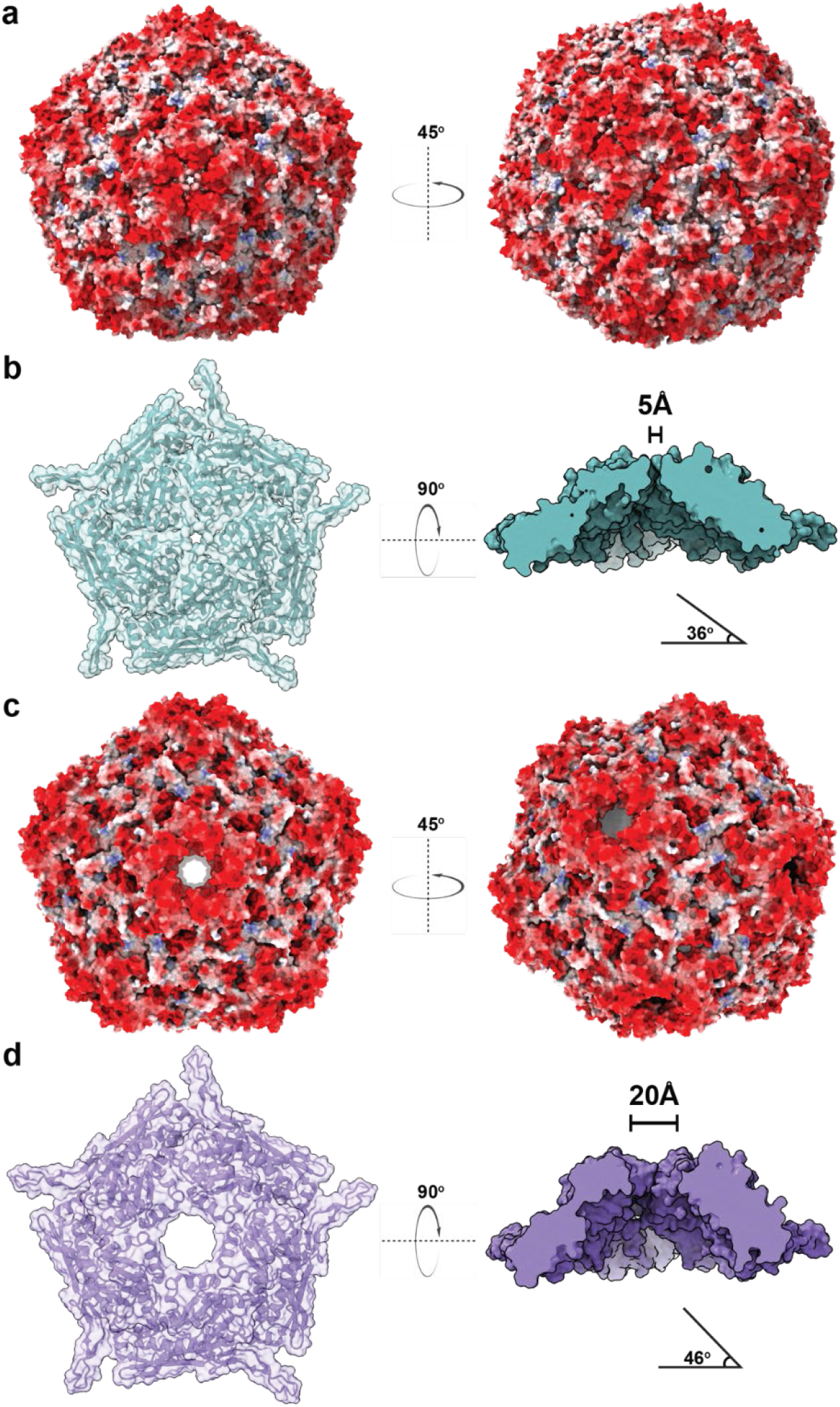
Structural overview of the AaEnc nanocage and 5-fold pore. **a**) Electrostatic surface representation of AaEnc in the “closed” conformation viewed down the 5-fold symmetry axis as well as at a 45° rightward turn (red, negative charge; white, neutral; blue, positive charge). **b**) Top-down ribbon and partially transparent surface representation of the “closed” AaEnc pentamer (left; cyan) and solid surface representation rotated 90° and viewed through the frontal axis plane to highlight pore size (right). The angle between protomers and the orthogonal of the 5-fold axis is highlighted. **c**) Electrostatic surface representation of AaEnc in the “open” conformation. **d**) Top-down ribbon and partially transparent surface representation of the “open” AaEnc pentamer (left; purple) and solid surface representation rotated 90° and viewed through the frontal axis plane to highlight pore size (right). The angle between protomers and the orthogonal of the 5-fold axis is highlighted.

Detailed examination of the “closed” and “open” state structures indicates that the primary conformational changes underlying the dynamic nature of the AaEnc pore are located at the apex of the so-called axial domain (A-domain) of the encapsulin protomer (**Figure 4a**). In addition, an overall backwards tilt of the “open” state protomer by 10° (**Figure 3b** and **Figure 3d**) also contributes to the observed size increase of the 5-fold pore (**Figure 3c**, **Figure 3d, Supplementary Video 2**). In the “closed” state, five pore residues – Asp185, His186, Gly187, Val188, and Pro189 – form a short loop between the α6 and α7 helices encompassing a predicted α-turn (**Figure S5**); whereas in the “open” state, the same residues form a tighter turn that loses the predicted α-turn conformation, with Val188 and Pro189 becoming part of and extending the α7 helix (**Figure 4c**, **Figure S5**, and **Figure S6**).^38^ His186 undergoes the most readily observable conformational change. It is located at the apex of the A-domain in the “closed” state, yet partially buried between two adjacent protomers in the “open” state (**Figure 4b**, **Figure 4c**). Furthermore, in the “open” state, intermolecular hydrogen bonding is observed between His186 and Asp150 as well as Ser181 and Asn157 of adjacent protomers (**Figure 4c**). These two hydrogen bonds are notably absent in the “closed” state. His186 is not strictly conserved among other structurally characterized Family 1 encapsulins (**Figure S1**) and cannot be used alone as an indicator for the presence of dynamic 5-fold pores in encapsulin shells, as in the *H. ochraceum* encapsulin, which also displays “closed” and “open” pore states, where His186 is substituted with Asp186 (**Figure S1** and **Figure S4**). Interestingly, in *H. ochraceum* the residue corresponding to Asp150 in AaEnc is Arg150. This could indicate that analogous to the hydrogen bonding between His186 and Asp150 (AaEnc), bonding between Asp186 and Arg150 (*H. ochraceum),* with swapped H-bond donors/acceptors, may be possible under certain conditions. However, this was not observed in the *H. ochraceum* “open” conformation.

**Figure 4.**
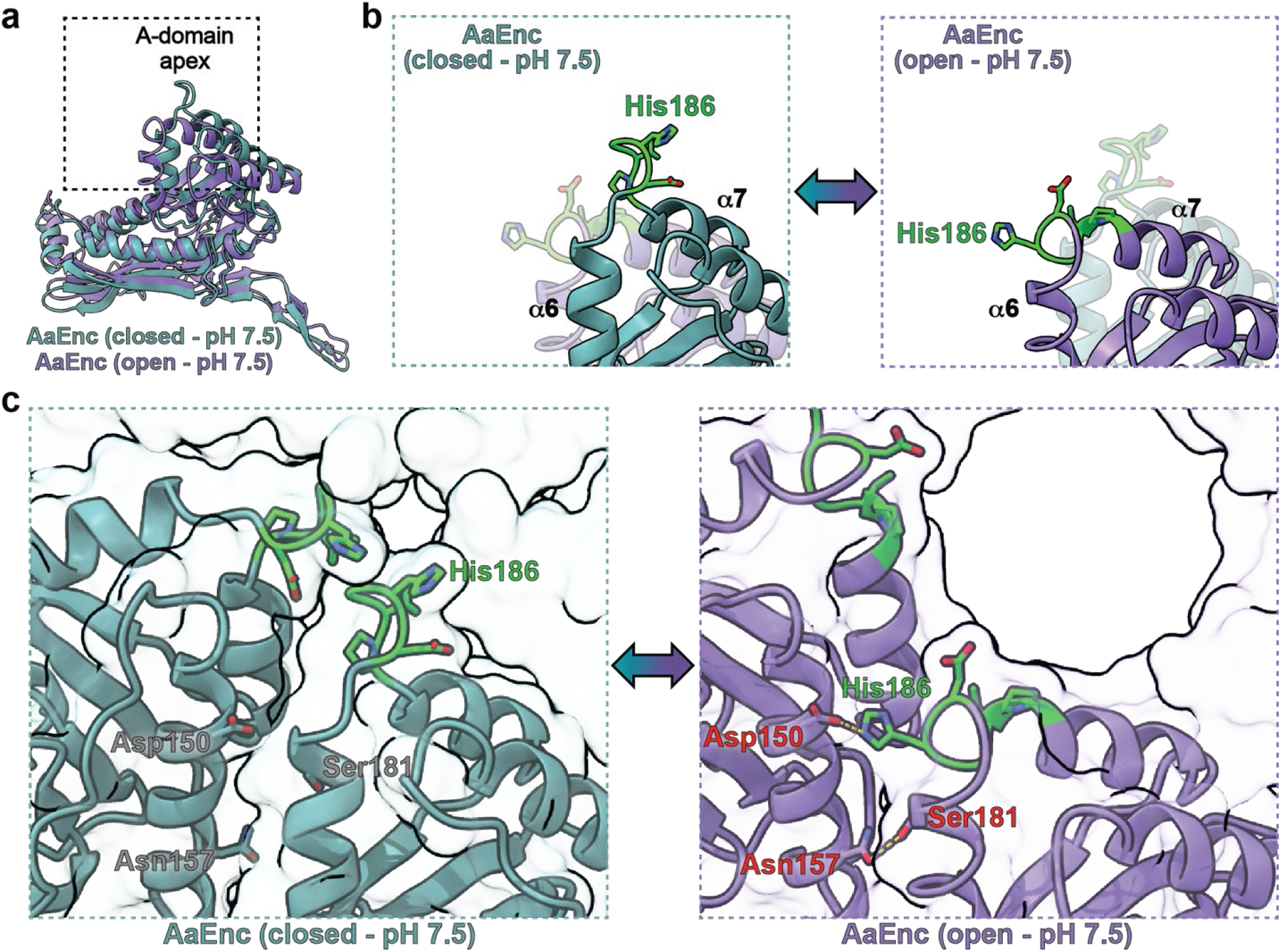
Detailed structural analysis of the AaEnc 5-fold pore. **a**) Aligned and overlayed ribbon representation of the “closed” (cyan) and “open” (purple) AaEnc protomers with dashed box highlighting the A-domain. **b**) Magnified ribbon representation juxtaposing the dynamic A-domain of the “closed” (cyan; left, solid; right, transparent) and “open” (purple; right, solid; left, transparent) AaEnc protomer, with the loop residues of interest–Asp185, His186 (labeled), Gly187, Val188, and Pro189– highlighted (green). **c**) Magnified solid ribbon representation of two adjacent AaEnc A-domains with transparent surface representation of the AaEnc pentamer highlighting the 5-fold pore. The “closed” state (left, cyan) exhibits a lack of hydrogen bonds between Asp150 (gray) and His186 (green) as well as Asn157 (gray) with Ser181 (gray), while the “open” state (right, purple) showcases gained hydrogen bonds between Asp150 (red) with His186 (green), as well as Asn157 (red) with Ser181 (red).

Overall, the cryo-EM density for the loop region of the *H. ochraceum* encapsulin in the “open” conformation was not well defined, whereas for AaEnc, both “closed” and “open” states exhibit strong and well-defined densities (**Figure S6**). This could in part be due to the additional stabilization of the “open” state in AaEnc resulting from the hydrogen bonding observed between His186 and Asp150.

Additional cryo-EM experiments were carried out at pH 3.0 to assess the structure of AaEnc under strongly acidic conditions. At pH 3.0, only a single “closed” conformational state was observed and was determined to 2.77 Å resolution (47,164 particles) (**Figure S7** and **Table S3**). It was also found that both of the “closed” AaEnc states – at pH 7.5 and pH 3.0 – are seemingly identical, with a root-mean-square deviation (RMSD) of 0.32 between the two aligned protomers (**Figure S8**).^39^

*In silico* analyses were conducted using the APBS-PDB2PQR software suite to quantitatively assess the hydrogen bonding and solvent exposure of the His186 and Asp150 residues at pH 7.5 and 3.0 (**Table 2**).^40^ The results further corroborate that His186 is more exposed – calculated as only 10% buried – in the “closed” states, and more buried – calculated as 39% buried – in the “open” state due to the inter-protomer hydrogen bonding described above. Furthermore, no hydrogen bonds were predicted between His186 and Asp150 in either of the “closed” states, while being clearly predicted for the “open” state.

Based on the results outlined above the “closed” state is clearly favored at low pH. It seems likely that in addition to global protonation state changes throughout the AaEnc protomer, specifically the protonation of Asp150 at low pH would preclude any hydrogen bonding with His186, thus making the conformational change from “closed” to “open” state energetically less favorable at acidic conditions. The specific molecular and biological functions of dynamic 5-fold pores in encapsulin shells is currently unknown. However, as the native cargo of AaEnc is a DyP-type peroxidase, generally known to be optimally active at acidic pH values, the preference of the AaEnc nanocage for the “closed” pore state at low pH might have significant functional and biological implications.

**Table 2.** *In silico* analysis of key pore residues. Calculated buriedness, hydrogen bonds, and pKa values are shown.

| State and residue | Buried | Sidechain Hydrogen Bond (Partner)<sup>1</sup> | Calculated pKa |
| --- | --- | --- | --- |
| **Closed (pH 7.5)** |  |  |  |
| Asp150 | 44% | -0.57 (Gln147, intramolecular) | 5.63 |
| His186 | 10% | none | 6.00 |
| **Open (pH 7.5)** |  |  |  |
| Asp150 | 12% | -0.53 (His186, intermolecular) | 3.50 |
| His186 | 39% | 0.53 (Asp150, intermolecular) | 6.63 |
| **Closed (pH 3.0)** |  |  |  |
| Asp150 | 43% | 0.90 (Asp149, intramolecular) | 7.64 |
| His186 | 10% | none | 5.85 |

<sup>1</sup> pKa shift due to respective hydrogen bond.

### Computational analysis of continuum electrostatics and solvation of the AaEnc protomer

To gain deeper insights into the stability of the AaEnc nanocage under acidic conditions, an array of computational analyses focused on continuum electrostatics and solvation of the AaEnc protomer was carried out. In particular, we thought to investigate if the AaEnc protomer would be predicted to show increased acid stability by itself outside the context of the encapsulin shell. The APBS-PDB2PQR software suite was used to assess the surface electrostatics of AaEnc at different pH values ranging from pH 2.0 to 8.0 (**Figure 5a**).^40^ The calculated protein electrostatics correlated well with the calculated pI of 4.41, showing a change from an overall positive surface charge below pH 4.0 to an overall negative surface charge above pH 5.0. Next, using the Protein-Sol software package, a heatmap depicting the average charge per residue based on pH and ionic strength was calculated.^41^ The results again correlate well with the calculated pI, with a positive average charge per residue at pH 4.0 and below, and a negative average charge per residue at pH 4.5 and above, regardless of ionic strength (**Figure 5b**). To investigate if a discernible increase in folded state protein stability mediated by interactions between ionizable groups might exist for the AaEnc protomer at acidic pH, further analyses using Protein-sol were carried out (**Figure 5c**). It was found that at physiological ionic strength (150 mM), per residue energies below pH 4.5 were positive, indicating decreased stability of the protomer fold below this pH threshold.

**Figure 5.**
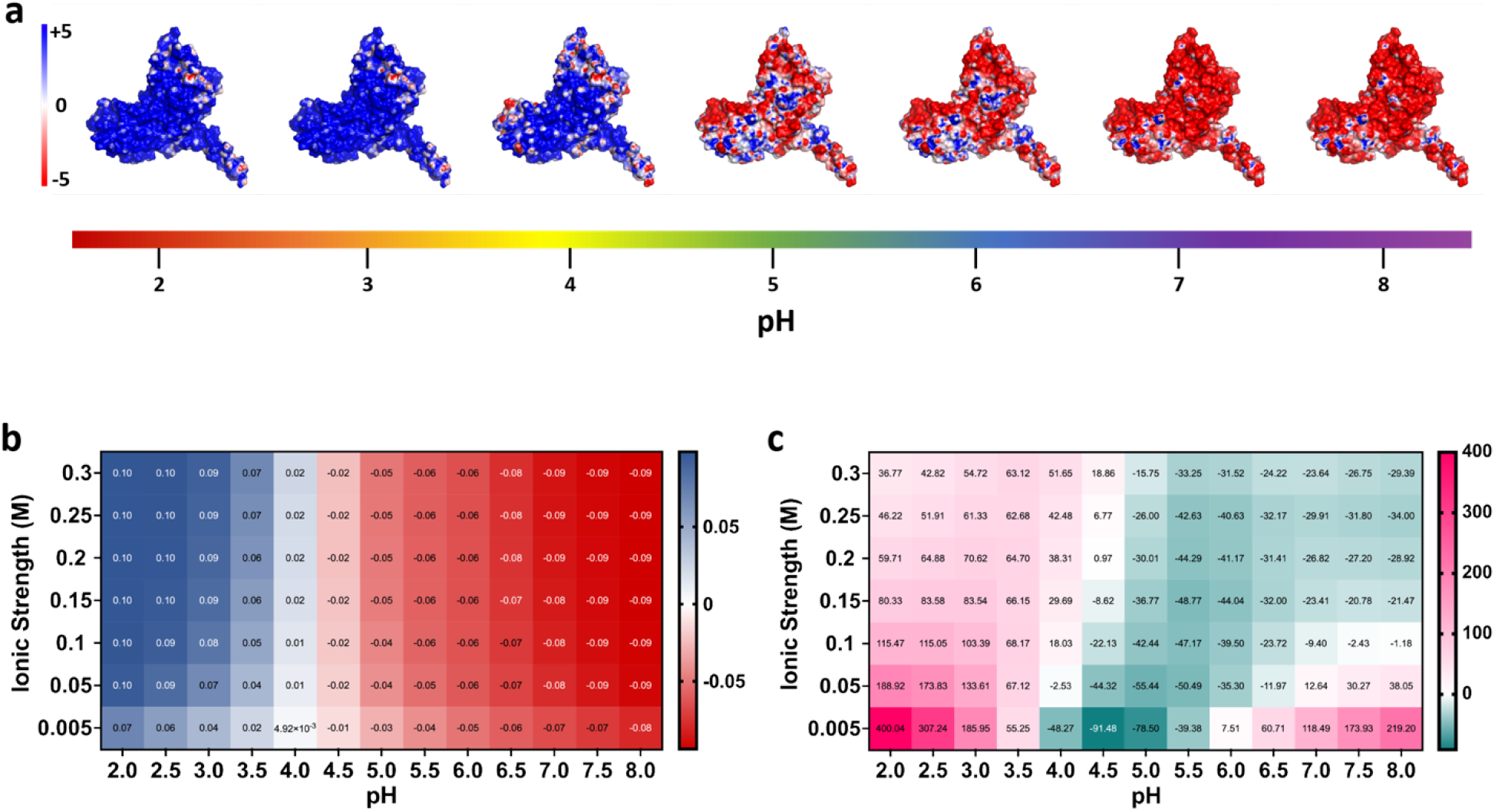
Computational characterization of the AaEnc protomer. **a**) Surface charge visualization of the AaEnc protomer with amino acid protonation states calculated by pH with pdb2pqr and protein electrostatics calculated with APBS. **b**) Heatmap of predicted average charge per residue for the AaEnc protomer at different pH and ionic strength values. **c**) Heatmap of predicted folded state protein stability as interactions between ionizable groups in joule per residue for the AaEnc protomer at different pH and ionic strength values.

Taken together, our computational analysis of the AaEnc protomer suggests that the observed acid stability of the AaEnc nanocage is not easily attributable to a highly stable protomer building block. Instead, AaEnc acid stability is likely due to a complex combination of factors at the scale of the assembled 60mer nanocage. Important factors likely include favorable inter-protomer interactions within the context of the encapsulin shell, such as the structural dynamics and hydrogen bonding interactions discussed above.

### *In vivo* cargo loading and pH effects on internalized cargo

To investigate if the AaEnc shell has any influence on the acid stability of internalized cargo proteins, heterologous cargo loading experiments were carried out followed by pH screens. The predicted C-terminal targeting peptide (TP) of the native AaEnc DyP cargo enzyme, was genetically fused to the C-terminus of eGFP (eGFP-TP) and cloned immediately upstream of the AaEnc gene for co-expression.^4,42^ eGFP was chosen as a non-native cargo due to its reliable expression, favorable solubility, simple detection, and predictable and well-reported pH sensitivity profile.^43,44^ *In vivo* eGFP-TP cargo loading was confirmed via its co-purification by SEC with co-expressed AaEnc, negative stain TEM analysis, and native polyacrylamide gel electrophoresis (PAGE) (**Figure 6a**, **Figure 6b**, and **Figure 6c**). Next, the fluorescence of equimolar amounts of free eGFP and AaEnc-encapsulated eGFP-TP were compared across a range of pH values from pH 3.0 to 8.0 using a plate-based fluorescence assay (**Figure 6d**). Both free eGFP and AaEnc-encapsulated eGFP-TP yielded very similar sigmoidal fluorescence response curves, demonstrating that the interior pH of AaEnc is not appreciably different from the bulk pH. Further, encapsulation within AaEnc does apparently not alter cargo pH sensitivity to a significant degree. These results indicate that the AaEnc shell does not represent an effective diffusion barrier for protons. Thus, buffer pH changes will result in the rapid equilibration of the external and luminal pH. Similar behavior has been observed for other protein-based compartments, particularly the carbon-fixing carboxysome bacterial microcompartment.^45^ Stopped-flow pH colorimetry indicated a rapid equilibration of the luminal carboxysome pH to that of the bulk solvent highlighting the porosity of the carboxysome shell towards protons. However, considering that one of the likely primary functions of encapsulin and carboxysome shells is to control the flux of specific small molecules into and out of the shell interior^45,46^ – possibly through the action of dynamic or gated pores – the idea that protein shells could be able to control the passage of protons is not out of the realm of possibility. A molecular mechanism similar to that employed by aquaporins, which allow passage of water molecules but block proton flux, could certainly exist or be engineered in protein shells.^47^

**Figure 6.**
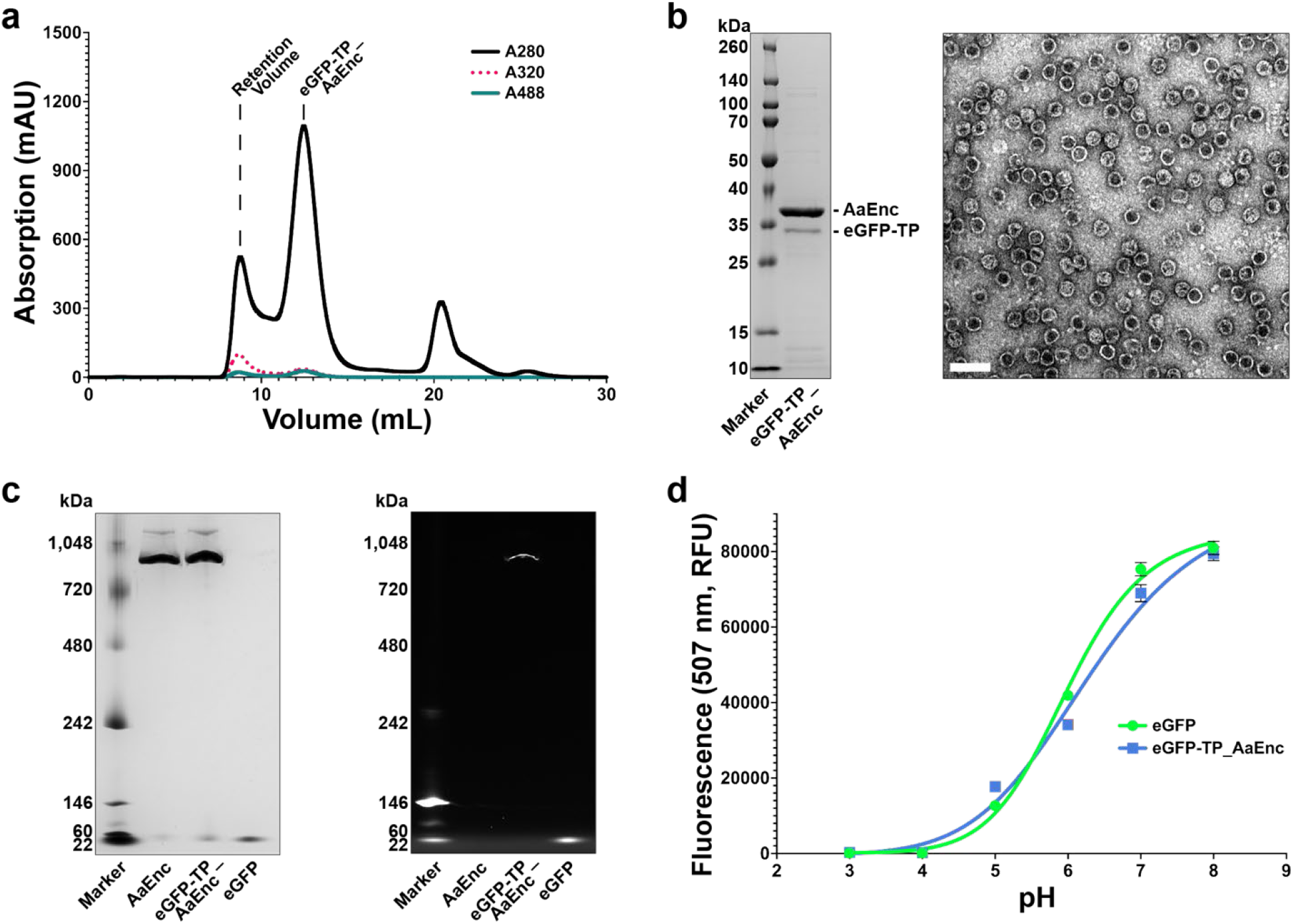
Analysis of eGFP cargo-loaded AaEnc. **a**) Size exclusion chromatography (SEC) analysis of AaEnc-encapsulated eGFP showing elution at 12-13 mL via protein absorbance at 280 nm and specific eGFP absorbance at 488 nm. **b**) SDS-PAGE analysis of purified eGFP-loaded AaEnc (left) and negative stain TEM (right). Scale bar: 50 nm. **c**) Native PAGE gel analysis of empty AaEnc, eGFP-loaded AaEnc, and free eGFP (left) along with corresponding fluorescence analysis of the same gel highlighting co-elution of eGFP fluorescence with the high molecular weight encapsulin band (right). **d**) Plate-based fluorescence analysis of AaEnc-encapsulated eGFP (eGFP-TP AaEnc) versus free eGFP. Data shown as means with error bars representing standard deviations from three independent experiments.

## Conclusion

The large number of encapsulin systems distributed across diverse bacterial and archaeal phyla – including many extremophiles – represents a largely untapped source of novel biotechnological tools.^4^ Ongoing discoveries and research within the encapsulin field has resulted in the characterization of many nano-encapsulation systems with a quickly expanding and diverse list of useful molecular features.^10,11,14,18,28^ However, relatively little attention has been focused on systematically exploring encapsulins from extremophilic bacteria and archaea with unusual molecular characteristics and stability profiles. With this study, we have taken the first step towards addressing this issue with a focus on the acid stability of the AaEnc nanocage.

Our results highlight the difficulty of pinpointing specific protein characteristics that lead to increased acid stability. Beyond the previously reported observation that acid stable proteins contain an increased number of aspartate and glutamate residues, resulting in a low pI,^48^ no other adaptations towards acid stability are readily apparent for AaEnc. We find that at the protomer level, AaEnc does not display any obvious properties beyond a low pI, that would indicate exceptionally high acid stability. This analysis is necessarily purely computational as encapsulin protomers quickly self-assemble to form protein nanocages and cannot be studied in isolation. It seems likely that the formation of a large 60mer protein complex plays a role in the acid tolerance of AaEnc with some assembled shells still present after extended incubation at pH 1.9. It can be speculated that minimizing the number of ionizable groups at key subunit interfaces within the AaEnc shell would contribute towards its stability at low pH. Beyond its unusual acid tolerance, AaEnc exhibits the unique feature of seemingly pH-dependent all “closed” or all “open” 5-fold pore states. At physiological pH, the “closed” and “open” states exist in a ratio of 2:1, whereas at low pH, the equilibrium is completely shifted towards the “closed” state. This behavior likely has important functional and biological implications and will require further study. Its stability, pH-responsive pores, and the fact that the “open” pore state with a diameter of 20 Å represents the largest encapsulin pore reported to date, make AaEnc an interesting target for future nanocage engineering applications in catalysis, nanotechnology, and medicine.

Specifically, acid-tolerant protein cages could offer novel opportunities in nanoreactor design and engineering, particularly in the context of industrial biopolymer degradation which requires acidic conditions. Enzyme encapsulation or co-localization could improve the performance of enzymes like chymosins,^49^ dye-decolorizing peroxidases,^50^ glucoamylases,^51^ and proteases,^52^ already extensively used in the food industry, agriculture, and biofuel production.^53–56^ Further, bioleaching and bioremediation approaches aimed at recovering valuable or toxic metals could benefit from acid stable nano-encapsulation systems able to sequester specific metal-binding enzymes of interest.^24,57^ Finally, a number of biomedical applications of protein nanocages related to drug delivery or intracellular targeting of acidic compartments could also benefit from robust and easily engineerable nanocages like _AaEnc._^58,59^

In sum, AaEnc is the first characterized highly acid stable encapsulin nanocage with a unique pH-dependent dynamic pore and, therefore, represents a novel useful tool for the nanocage engineering community.

## Methods

### Chemicals and biological materials

All chemicals were used as supplied by vendors without further purification. Imidazole, Invitrogen Novex WedgeWell 14% tris-glycine Mini Protein Gels, Isopropy-ß-D-thiogalactopyranoside (IPTG), lysozyme, NativePAGE™ 4 to 16% bis-tris Mini Protein Gels, NativeMark Unstained Protein Standard, Spectra™ Multicolor Broad Range Protein Ladder, Thermo Scientific Pierce 660 nm Protein Assay Reagent, Tris base, Tris HCl, all restriction enzymes, and all cell culture media and reagents were purchased from Fisher Scientific, Inc. (USA). Gibson Assembly Master Mix was purchased from NEB (USA). Amicon Ultra-0.5 mL centrifugal units and Benzonase^®^ nuclease were purchased from MilliporeSigma (USA). BL21 (DE3) Electrocompetent Cells used for *E. coli* expression were also purchased from MilliporeSigma (USA). Bis-tris propane from Research Products International (USA) was used for the assembly buffer. Ni-NTA agarose from Gold Biotechnology, Inc. (USA) was used for His-tagged protein purification.

### Instrumentation

Cell lysis was conducted via sonication with a Model 120 Sonic Dismembrator from Fisher Scientific, Inc. (USA). Protein was quantified on a Nanodrop Spectrophotometer from ThermoFisher Scientific, Inc. (USA). Protein purification was carried out on an AKTA Pure fast liquid protein chromatography system; size exclusion chromatography (SEC) was carried out with a HiPrep 16/60 Sephacryl S-500 HR and Superose 6 10/300 GL columns (Cytiva, USA); anion exchange was carried out with a HiTrap Q FF column (Cytiva, USA). Polyacrylamide gel electrophoresis (PAGE) and NativePAGE were performed in an XCell SureLock from Invitrogen/ThermoFisher Scientific (USA). Gel images were captured using a ChemiDoc Imaging System from Bio-Rad Laboratories, Inc. (USA). DLS was carried out on an Uncle from Unchained Labs (USA). TEM was carried out on a Morgagni 100 keV Transmission Electron Microscope (FEI, USA). Plate-based fluorescence assays were conducted on the Synergy H1 Microplate Reader from BioTek Instruments (USA). EM grid glow discharging was conducted with a PELCO easiGlow™ system by Ted Pella, Inc (USA). A Talos Arctica Cryo Transmission Electron Microscope by ThermoScientific, Inc. (USA) equipped with a K2 Summit direct electron detector by Gatan, Inc. (USA) located at the University of Michigan Life Sciences Institute was used for cryo-EM. Smaller materials are listed along with corresponding methods below.

### Software

The following software was used throughout this work: Adobe Illustrator 2021 v25.0.0 (figures), cryoSPARC v3.3.1^60^ (cryo electron microscopy), Fiji/ImageJ v2.1.0/1.53c^61^ (densitometric data analysis and TEM images), GraphPad Prism for Mac OS v9.4.0 (chromatography, melting temperature/aggregation, and fluorescence graphs), Bio-Rad Image Lab Touch Software (gel imaging), Microsoft Excel for Mac v16.46 (DLS graphs), Phenix v1.19.2-4158^62^ (model building), UCSF Chimera v1.16^63^ and ChimeraX v3^39^ (cryo-EM density and model visualization), and UNICORN 7 (FPLC system control and chromatography). Online software suites or tools are listed along with corresponding methods below.

### Bioinformatic search for acid-stable encapsulins

A curated list of Family 1 encapsulins^19^ was sorted according to molecular weight, removing any entries falling below 25 kDa or above 35 kDa to remove partial annotations and fusion encapsulins, respectively. Results were then processed via the Expasy Compute pI/MW tool (https://web.expasy.org/compute_pi/). Data was then organized according to calculated pI and binned for histogram analysis or plotted for scatterplot analysis via GraphPad Prism.

### Sequence alignments

Encapsulin alignments were generated with the ESPript 3 server (http://espript.ibcp.fr/) using a protein sequence alignment produced with Clustal Omega, with secondary structure information based on the TmEnc structure (PDB 3DKT; **Figure S1**) or the “open” and “closed” AaEnc structures (**Figure S5**).^38,64^

### Protein production

For all target proteins, plasmids were constructed with target *E. coli* codon-optimized gBlock genes, synthesized by IDT (USA), inserted into the pETDuet-1 vector via Gibson assembly using the NdeI and PacI restriction sites (**Table S2)**. *E. coli* BL21 (DE3) was transformed with the respective plasmids via electroporation per protocol and 25% glycerol bacterial stocks were made and stored at −80°C until further use. Starter cultures were grown in 5 mL LB with 100 mg/mL ampicillin at 37°C overnight. For all constructs, 500 mL of LB with ampicillin was inoculated with overnight starter cultures and grown at 37°C to an OD600 of 0.4-0.5, then induced with 0.1 mM IPTG and grown further at 30°C overnight for ca. 18 h. Cells were then harvested via centrifugation at 10,000 rcf for 15 minutes at 4°C and pellets were frozen and stored at −80°C until further use.

### Protein purification

Frozen bacterial pellets were thawed on ice and resuspended in 5 mL/g (wet cell mass) of cold Tris Buffered Saline (20 mM Tris pH 7.5, 150 mM NaCl). Lysis components were added (0.5 mg/mL lysozyme, 1 mM tris(2-carboxyethyl)phosphine [TCEP], one SIGMA*FAST* EDTA-free protease inhibitor cocktail tablet per 100 mL, 0.5 mM MgCl2, and 25 units/mL Benzonase^®^ nuclease) and samples were lysed on ice for 10 min. Samples were then sonicated at 60% amplitude for 5 min total (eight seconds on, 16 seconds off) until no longer viscous. After sonication, samples were centrifuged at 8,000 rcf for 15 minutes at 4°C. Samples were then subjugated to 10% polyethylene glycol (PEG) 8000 precipitation (lysate brought to 10% PEG 8K and 500 mM NaCl and incubated for 30 minutes on ice, then centrifuged 8,000 rcf for 15 min). Supernatant was discarded and the pellet was resuspended in 5 mL TBS pH 7.5 and filtered using a 0.22 μm syringe filter (Corning, USA). The protein sample was then loaded on an AKTA Pure and purified via a Sephacryl S-500 column. Sample fractions were pooled and buffer exchanged into AIEX Buffer (20 mM Tris pH 7.5) and loaded onto an AKTA Pure, then purified via HiTrap Q-FF by linear gradient into AIEX Buffer with 1M NaCl. Sample flow-through was collected and centrifuged at 10,000 rcf for 10 min, then loaded on an AKTA Pure for final purification via a Superose 6 10/300 GL column pre-equilibrated with TBS pH 7.5. All proteins were stored at 4°C until use.

For free His-tagged eGFP purification, the sample was lysed as above in NTA Resuspension Buffer (50 mM Tris pH 8.0, 150 mM NaCl, 1 mM TCEP, 10 mM imidazole, and 5% glycerol). Lysate was bound to Ni-NTA resin pre-equilibrated with NTA Resuspension Buffer via rocking at 4°C for 45 minutes. Supernatant was discarded and the bound sample was washed once with NTA Resuspension Buffer and a second time with NTA Resuspension Buffer with 20 mM imidazole. Free His-tagged eGFP was then eluted three times with Elution Buffer (50 mM Tris pH 8.0, 150 mM NaCl, 1 mM TCEP, 350 mM imidazole, 5% glycerol) and stored at 4°C for future use.

### Transmission electron microscopy

Samples were diluted to 0.1-0.3 mg/mL and buffer exchanged into various buffers ranging from pH 1.5 to pH 10.0 (**Figure 2c** and **Figure S2**) via five successive exchanges in 100 kDa MWCO Amicon Ultra-0.5 mL centrifugal units. Buffers used consisted of 50 mM sodium phosphate and 150 mM NaCl, pH 1.5; 50 mM sodium phosphate and 150 mM NaCl, pH 1.9; 50 mM sodium phosphate and 150 mM NaCl, pH 2.1; 50 mM sodium phosphate and 150 mM NaCl, pH 2.25; 50 mM sodium phosphate and 150 mM NaCl, pH 3.0; 50 mM sodium citrate and 150 mM NaCl, pH 5.0; 50 mM MES and 150 mM NaCl, pH 6.0; 50 mM bistris propane and 150 mM NaCl, pH 9.0; and 50 mM CHES and 150 mM NaCl, pH 10.0. Samples were incubated at 4°C for six hours, and then immediately stained and imaged. Additional pH 2.25 and pH 3.0 samples were stored at 4°C for two days and stained and imaged. Negative stain transmission electron microscopy (TEM) was carried out on the various samples with 200-mesh gold grids coated with extra thick (25-50 nm) formvar-carbon film (EMS, USA) made hydrophilic by glow discharging at 5 mA for 60 s. Briefly, 3.5 μL of sample was added to the grid and incubated for 30 seconds, wicked with filter paper, and washed once with distilled water and once with 0.75% (w/v) uranyl formate before staining with 8.5 μL of uranyl formate for 30 seconds. TEM images were captured using a Morgagni transmission electron microscope at 100 keV at the University of Michigan Life Sciences Institute. For all TEM experiments, samples were roughly 0.2 mg/mL of AaEnc monomer in appropriate buffer.

### Dynamic and static light scattering analyses

All sizing and polydispersity measurements were carried out on an Uncle by Unchained Labs (USA) at 30 °C in triplicate. Purified AaEnc samples were adjusted to 0.4 mg/mL of monomer in the appropriate corresponding buffers and centrifuged at 10,000 rcf for 10 min, then immediately analyzed via DLS (**Figure 2b**). Static light scattering aggregation temperature (T_agg_) analysis was then conducted on similarly prepared samples over a 25°C to 95°C ramp at 1°C per minute (**Figure 2d**).

### Protease assays

AaEnc and bovine serum albumin (BSA; ThermoScientific Pierce, USA) were individually buffer exchanged into Pepsin Assay Buffer (50 mM Na2PO4, 150 mM NaCl, pH 3.0) and mixed in a 40:1 molar ratio with commercially purchased pepsin protease (Promega, USA), then incubated at 37°C for 3 h and frozen until later use. Purified AaEnc and BSA were individually buffer exchanged into TBS pH 7.5 and mixed in a 40:1:1 molar ratio with commercially purchased trypsin (Promega, USA) and chymotrypsin (Promega, USA), then incubated at 37°C for 8 h and frozen until later use. All samples were then rapidly thawed and examined via PAGE analysis.

### Cryo-electron microscopy

#### Sample preparation

The purified protein samples were concentrated to 3 mg/mL in 150 mM NaCl, 20 mM Tris pH 7.5 or 150 mM NaCl, 50 mM Na2PO4 pH 3.0. 3.5 μL of protein samples were applied to freshly glow discharged Quantifoil R1.2/1.3 grids and plunged into liquid ethane using an FEI Vitrobot Mark IV (100% humidity, 22°C, blot force 20, blot time 4 seconds, drain time 0 seconds, wait time 0 seconds). The frozen grids were clipped and stored in liquid nitrogen until data collection.

#### Data collection

Cryo-electron microscopy movies were collected using a ThermoFisher Scientific Talos Arctica operating at 200 keV equipped with a Gatan K2 Summit direct electron detector. Movies were collected at 45,000x magnification using the Leginon^65^ software package with a pixel size of 0.91 Å/pixel and an exposure time of 5 or 8 s, frame time of 200 ms, and total dose of 42 e^-^/A^2^ for the pH 3.0 sample and 41 e^-^/A^2^ for the pH 7.5 sample. 1,357 movies were collected for the pH 3 sample and 975 movies were collected for the pH 7.5 sample.

#### Data processing

pH 7.5 sample: All data processing was performed using cryoSPARC v3.3.1.^60^ 975 Movies were imported and motion corrected using Patch Motion Correction and CTF fits were refined using Patch CTF. 821 movies with CTF fits better than 8.0 Å were selected for downstream processing. Roughly 200 particles were picked manually using Manual Picker and grouped into 10 classes using 2D Classification. Well resolved classes were selected and used as templates for Template Picker to pick particles with a specified particle diameter of 240 Å. 56,583 particles with a box size of 384 pixels were extracted and subjected to 3 rounds of 2D Classification with 100 classes yielding 44,686 particles in good classes. Ab-Initio Reconstruction with 6 classes and I symmetry was carried out next. The two main classes were selected representing the all “closed” (29,056 particles) and all “open” (13,581 particles) states. Particles from each respective state were used as inputs for separate Homogenous Refinement jobs (with I or C1 symmetry) with the following settings: optimize per-particle defocus, optimize per-group CTF params, and Ewald Sphere correction enabled. The I refinements yielded a 2.90 Å density for the “closed” state, and a 3.32 Å density for the “open” state, whereas the C1 refinements resulted in 4.84 Å and 4.44 Å maps, respectively (**Figure S3**). 3D Variability Analysis with 3 components was carried out on both particle sets using the C1 Refinement results as inputs, however, no components could be resolved corresponding to “open” and “closed” states within the same density.

pH 3.0 sample: The same preprocessing procedure was used as for the pH 7.5 sample yielding 52,210 extracted particles with a box size of 384 pixels. 3 rounds of 2D classification with 100 classes resulted in 47,599 good particles. Ab-Initio Reconstruction with 6 classes and I symmetry was carried out yielding one dominant class containing 47,164 particles. This was followed by I and C1 Homogenous Refinement jobs using the following parameters: optimized per-particle defocus, optimize per-group CTF params, and Ewald Sphere correction. The I refinement resulted in a density of 2.77 Å while the C1 refinement yielded a 4.12 Å map (**Figure S7**).

#### Model building

A homology model was generated using RoseTTAFold^66^ on the Robetta server and was used as a starting model for all model building efforts. This starting model was manually placed into the respective cryo-EM maps using Chimera v1.16,^63^ and was further fit using the Fit to Volume command. The placed monomeric models were then manually refined against the respective cryo-EM maps using Coot v8.9.6.^67^ The resulting models were further refined using Real Space Refine in Phenix v 1.19.2-4158^68^ with default settings and three iterations. After inspecting the refined models in Coot, symmetry restraints were pulled from the maps using the Phenix.Find_NCS_from_Map command with I symmetry. Complete shell models were assembled using the Phenix.Build_from_NCS command. These shell models were then used as inputs for a final round of Real Space Refine with NCS restraints, 3 iterations, and all other settings set to default. The models were deposited to the PDB under PDB ID 8DN9, 8DNL, and 8DNA; and the EMDB under EMD-27558, EMD-27573, and EMD-27560.

### Computational electrostatics and solvation analyses

*In silico* hydrogen bonding and buriedness analyses were conducted using the APBS-PDB2PQR software suite (https://server.poissonboltzmann.org/pdb2pqr) with PROPKA v3.2 to predict pKa values and assign protonation states at the provided pH values. Analyses were conducted using the “closed” and “open” state AaEnc structures at pH 7.5 as well as the “closed” AaEnc structure at pH 3.0. The AaEnc protomer was further analyzed using APBS-PDB2PQR to assess the calculated surface charge of AaEnc across various pH values from pH 2.0 to pH 8.0 (**Figure 5a**).^40^ Calculated protomer charge and stability heatmaps were generated with the Protein-Sol webtool (https://proteinsol.manchester.ac.uk/heatmap) using the AaEnc protomer from the “closed” state at pH 7.5 as the input (**Figure 5b** and **Figure 5c**).^41^ Monomer structure alignments were carried out in ChimeraX.^39^

### *In vivo* cargo loading, native PAGE, and fluorescence analysis

The AaEnc encapsulated eGFP-TP sample was co-expressed, purified, and analyzed via TEM in the same manner as AaEnc as described above (**Figure 6a and Figure 6b**). The free His-tagged eGFP and the AaEnc-encapsulated eGFP-TP samples were concentrated to equimolar concentrations as determined by densitometric analysis via SDS-PAGE using Fiji/ImageJ.^61^ Empty AaEnc and eGFP-TP-loaded AaEnc were similarly concentrated to equimolar concentrations for comparative NativePAGE analysis.

All NativePAGE analyses were conducted in an Invitrogen XCell SureLock using NativePAGE™ 4 to 16% bis-tris mini protein gels and NativeMark Unstained Protein Standard from Fisher Scientific (USA) with 1x running buffer made from 10x Tris/Glycine Buffer from Bio-Rad Laboratories, Inc. (USA). 20 μg of protein was loaded per well, with effort to maintain equivalent amounts across all lanes for comparative analysis. NativePAGE gels were run overnight at 65 V for 16.5 hours at 4°C. The following day, gels were imaged via fluorescence imaging on a ChemiDoc Imaging System by Bio-Rad Laboratories, Inc. (USA), then stained with ReadyBlue™ Protein Gel Stain from Sigma-Aldrich (USA) and imaged and analyzed.

His-tagged eGFP and AaEnc-encapsulated eGFP-TP were buffer exchanged into varying pH buffers from pH 3.0 to pH 8.0 via five successive exchanges in 100 kDa MWCO Amicon Ultra-0.5 mL centrifugal units and incubated at 22°C for three hours. Endpoint eGFP fluorescence (488 nm/507 nm) was then measured in a BioTek Synergy H1 microplate reader at a final volume of 100 μL in Corning^®^ 96-well flat clear bottom black polystyrene microplates (**Figure 6c**).

## Acknowledgements

We gratefully acknowledge funding from the NIH (R35GM133325). Research reported in this publication was supported by the University of Michigan Cryo-EM Facility (U-M Cryo-EM). U-M Cryo-EM is grateful for support from the U-M Life Sciences Institute and the U-M Biosciences Initiative. Molecular graphics and analyses performed with UCSF ChimeraX, developed by the Resource for Biocomputing, Visualization, and Informatics at the University of California, San Francisco, with support from National Institutes of Health R01GM129325 and the Office of Cyber Infrastructure and Computational Biology, National Institute of Allergy and Infectious Diseases.

## Author Contributions

J.A.J. and T.W.G. designed the project. J.A.J. conducted the laboratory experiments and negative stain transmission electron microscopy, while M.P.A. collected and analyzed cryo-EM data. J.A.J. and M.P.A. built the AaEnc structural models. J.A.J. wrote the manuscript. T.W.G. processed cryo-EM data, edited the manuscript and oversaw the project in its entirety.

## Competing Interests

The authors declare no competing interests.

## Additional Information

Correspondence and requests for materials should be addressed to T.W.G.

## Notes

### Competing Interest Statement

The authors have declared no competing interest.

### Summary of Updates

Spelling mistake was corrected: Acidopropionibacterium acidopropionici was changed to Acidipropionibacterium acidipropionici.

