## Supplementary Information for "Exploring the Extreme Acid Tolerance of a Dynamic Protein Nanocage"

### 1. Comparative charge compositions of Family 1 encapsulins

**Table S1.** Charge composition of Family 1 encapsulins in the two lowest pI bins by organism and pI.

| Organism (UniProt ID) | pI[1] | # Acidic residues | # Basic residues | Acid/base ratio | Charge at pH 7.0[2] |
|---|---|---|---|---|---|
| *Halarsenatibacter silvermanii*[3] (A0A1G9H1B5_9FIRM) | 4.14 | 61 | 24 | 2.54 | -38.96 |
| *Desulfobacterales bacterium* (A0A1E7H417_9DELT) | 4.32 | 37 | 20 | 1.85 | -20.80 |
| *Halanaerobium sp. T82-1* (A0A139D0R2_9FIRM) | 4.35 | 54 | 28 | 1.93 | -26.04 |
| *Halanaerobium congolense* (A0A1G6LYG6_9FIRM) | 4.35 | 54 | 28 | 1.93 | -26.04 |
| *Halanaerobium congolense* (A0A318E778_9FIRM) | 4.35 | 54 | 28 | 1.93 | -26.04 |
| *Halanaerobium sp* (A0A315R6B2_9FIRM) | 4.36 | 53 | 28 | 1.89 | -25.04 |
| *Halanaerobium saccharolyticum* (A0A2T5RKL7_9FIRM) | 4.36 | 53 | 26 | 2.04 | -25.04 |
| *Halanaerobium kushneri* (A0A1N6TPV1_9FIRM) | 4.38 | 54 | 29 | 1.86 | -25.04 |
| *Halanaerobium sp. ST460_2HS_T2* (A0A368WL76_9FIRM) | 4.38 | 54 | 29 | 1.86 | -25.04 |
| *Acidipropionibacterium virtanenii* (A0A344UW06_9ACTN) | 4.39 | 43 | 26 | 1.65 | -25.37 |
| *Halanaerobium sp* (A0A315R3Q5_9FIRM) | 4.40 | 51 | 28 | 1.82 | -23.04 |
| *Halanaerobium saccharolyticum* (A0A4R6SET4_9FIRM) | 4.40 | 51 | 28 | 1.82 | -23.04 |
| *Acidipropionibacterium acidipropionici*[4] (K7RV67_ACIA4) | 4.41 | 41 | 25 | 1.64 | -23.47 |
| *Halanaerobium congolense* (A0A1H9Y5I8_9FIRM) | 4.41 | 53 | 29 | 1.83 | -24.04 |
| *Acidipropionibacterium acidipropionici* (A0A3Q9CQJ7_9ACTN) | 4.42 | 41 | 25 | 1.64 | -23.47 |
| *Tessaracoccus bendigoensis* (A0A1M6IV29_9ACTN) | 4.44 | 40 | 24 | 1.67 | -21.53 |
| *Halanaerobium saccharolyticum* (A0A4R6M0G8_9FIRM) | 4.45 | 52 | 30 | 1.73 | -22.04 |
| *Synergistaceae bacterium* (A0A3C0C467_9BACT) | 4.45 | 43 | 25 | 1.72 | -19.16 |
| *Actinomyces radicidenti* (A0A0X8JCJ6_ACTRD) | 4.46 | 52 | 31 | 1.68 | -26.45 |
| *Orenia metallireducens* (A0A285HJ09_9FIRM) | 4.46 | 48 | 28 | 1.71 | -21.85 |
| *Tsukamurella pseudospumae* (A0A138A052_9ACTN) | 4.47 | 40 | 25 | 1.60 | -17.77 |
| *Halanaerobium saccharolyticum* (A0A4R7Z6A4_9FIRM) | 4.47 | 52 | 30 | 1.73 | -22.04 |
| *Cloacibacillus porcorum* (A0A1B2I695_9BACT) | 4.47 | 44 | 27 | 1.63 | -18.16 |
| *Luteipulveratus mongoliensis* (A0A0K1JIL3_9MICO) | 4.49 | 42 | 27 | 1.56 | -22.27 |
| *Mycolicibacillus trivialis* (A0A1X2EII3_9MYCO) | 4.49 | 43 | 28 | 1.54 | -18.73 |
| *Halanaerobium praevalens* (E3DPB0_HALPG) | 4.49 | 51 | 30 | 1.70 | -21.04 |
| *Halanaerobium salsuginis* (A0A1I4N6V9_9FIRM) | 4.49 | 52 | 31 | 1.68 | -21.04 |
| *Halanaerobium hydrogeniformans* (E4RN20_HALHG) | 4.49 | 50 | 30 | 1.67 | -20.95 |
| *Orenia metallireducens* (A0A1C0A753_9FIRM) | 4.49 | 48 | 29 | 1.66 | -21.75 |

[1] Theoretical isoelectric point calculated with Expasy Compute pI/MW tool (https://web.expasy.org/compute_pi/).[8]
[2] Charge calculated via Geneious Prime 2020.2.4 (https://www.geneious.com).
[3] Single representative from lowest pI bin centered at 4.2 (*H. silvermanii*).
[4] AaEnc encapsulin data (red).

### 2. Sequence alignment of characterized encapsulin proteins

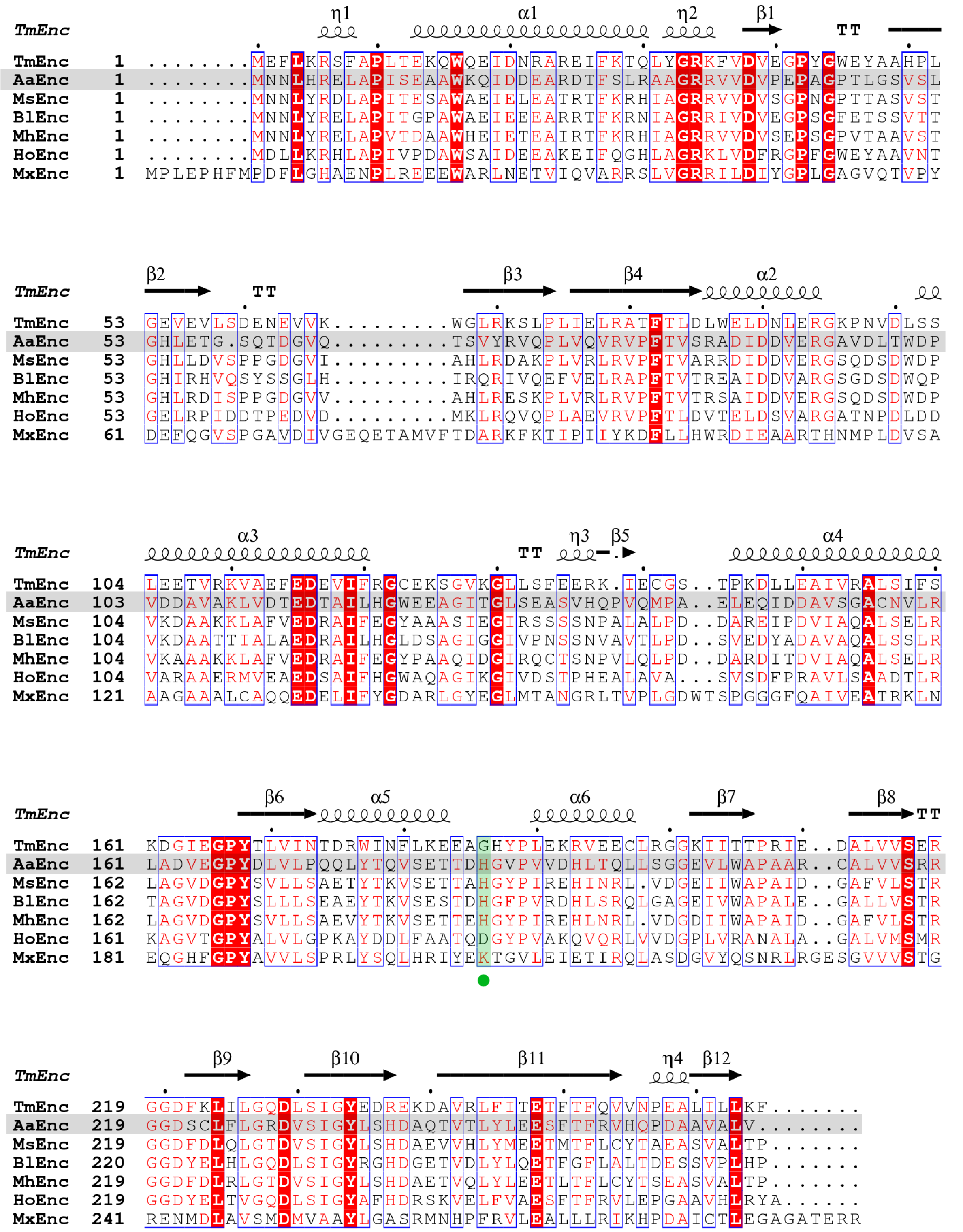

**Figure S1.** Sequence alignment of AaEnc to well characterized encapsulin proteins. Sequences aligned to *Thermotoga maritima* (TmEnc; PDB 3DKT),[1] including *Acidipropionibacterium acidipropionici* (ATCC 4875; AaEnc), *Mycobacterium smegmatis* (MsEnc; PDB 7BOJ),[2] *Brevibacterium linens* (*BlEnc;* UniProt ID A0A0B8ZXE8*)*,[3] *Mycolicibacterium hassiacum* (MhEnc; PDB 6I9G),[4] *Haliangium ochraceum* (PDB 7ODW)[5], and *Myxococcus xanthus* (PDB 4PT2).[6] The AaEnc sequence is highlighted in grey. Identical aligned residues are highlighted in red, with residues aligned to the His186 residue of interest from AaEnc highlighted in green and marked by a green circle. Alignment generated with the ESPript 3 server (http://espript.ibcp.fr/).[7]

### 3. Protein sequence data

**Table S2**. Protein sequences of constructs used.

| Construct | Protein sequence[1,2] |
|---|---|
| AaEnc | MNNLHRELAPISEAAWKQIDDEARDTFSLRAAGRRVVDVPEPAGPTLGSVSLGHLETG SQTDGVQTSVYRVQPLVQVRVPFTVSRADIDDVERGAVDLTWDPVDDAVAKLVDTED TAILHGWEEAGITGLSEASVHQPVQMPAELEQIDDAVSGACNVLRLADVEGPYDLVLP QQLYTQVSETTDHGVPVVDHLTQLLSGGEVLWAPAARCALVVSRRGGDSCLFLGRDVS IGYLSHDAQTVTLYLEESFTFRVHQPDAAVALV* |
| eGFP | MGSSHHHHHHGGSGMVSKGEELFTGVVPILVELDGDVNGHKFSVSGEGEGDATYGKL TLKFICTTGKLPVPWPTLVTTLTYGVQCFSRYPDHMKQHDFFKSAMPEGYVQERTIFFK DDGNYKTRAEVKFEGDTLVNRIELKGIDFKEDGNILGHKLEYNYNSHNVYIMADKQKN GIKVNFKIRHNIEDGSVQLADHYQQNTPIGDGPVLLPDNHYLSTQSALSKDPNEKRDH MVLLEFVTAAGITLGMDELYK* |
| eGFP-TP_AaEnc | MVSKGEELFTGVVPILVELDGDVNGHKFSVSGEGEGDATYGKLTLKFICTTGKLPVPWP TLVTTLTYGVQCFSRYPDHMKQHDFFKSAMPEGYVQERTIFFKDDGNYKTRAEVKFEG DTLVNRIELKGIDFKEDGNILGHKLEYNYNSHNVYIMADKQKNGIKVNFKIRHNIEDGSV QLADHYQQNTPIGDGPVLLPDNHYLSTQSALSKDPNEKRDHMVLLEFVTAAGITLGMD ELYKGGSGGSGGSGGPRDDSLGVGGLRGTPQLGSIPTPGRT*…MNNLHRELAPISEAA WKQIDDEARDTFSLRAAGRRVVDVPEPAGPTLGSVSLGHLETGSQTDGVQTSVYRVQP LVQVRVPFTVSRADIDDVERGAVDLTWDPVDDAVAKLVDTEDTAILHGWEEAGITGLS EASVHQPVQMPAELEQIDDAVSGACNVLRLADVEGPYDLVLPQQLYTQVSETTDHGV PVVDHLTQLLSGGEVLWAPAARCALVVSRRGGDSCLFLGRDVSIGYLSHDAQTVTLYLE ESFTFRVHQPDAAVALV* |

[1] Intergenic sequences for encapsulated constructs represented by ellipses.

[2] Stops indicated by asterisks.

### 4. Transmission electron microscopy analysis of AaEnc

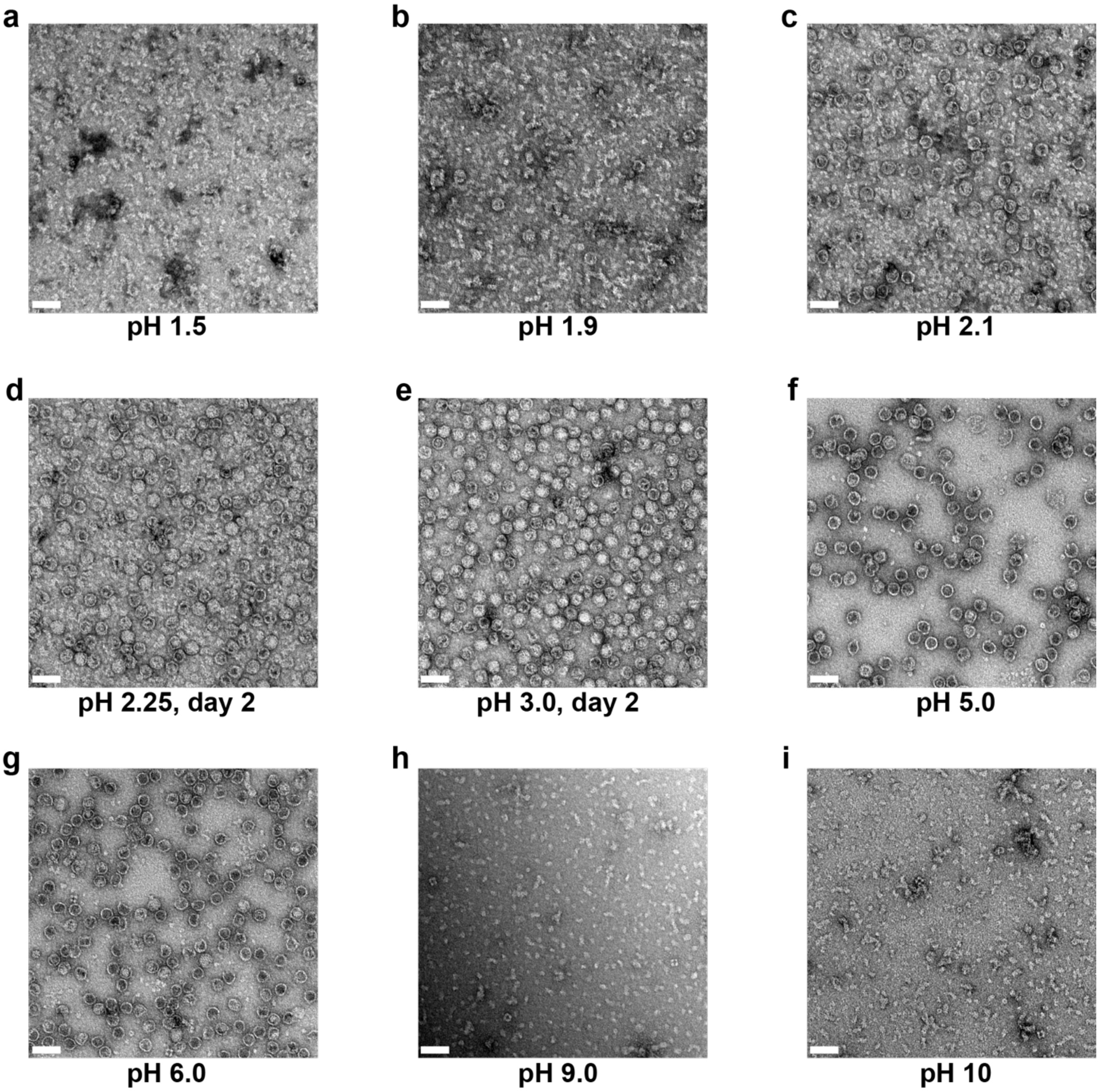

**Figure S2**. Additional TEM micrographs of the AaEnc nanocage. a) AaEnc in 50 mM phosphate buffer and 150 mM NaCl, pH 1.5. b) AaEnc in 50 mM phosphate buffer and 150 mM NaCl, pH 1.9. c) AaEnc in 50 mM phosphate buffer and 150 mM NaCl, pH 2.1. d) AaEnc in 50 mM phosphate buffer and 150 mM NaCl, pH 2.25 after two days of incubation at 4°C. e) AaEnc in 50 mM phosphate buffer and 150 mM NaCl, pH 3.0 after two days of incubation at 4°C. f) AaEnc in 50 mM sodium citrate buffer and 150 mM NaCl, pH 5.0. g) AaEnc in 50 mM MES buffer and 150 mM NaCl, pH 6.0. h) AaEnc in 50 mM Bis-tris propane buffer and 150 mM NaCl, pH 9.0. i) AaEnc in 50 mM CHES buffer and 150 mM NaCl, pH 10.0. All samples were buffer exchanged and incubated for six hours unless otherwise stated. Scale bars (white): 50 nm.

### 5. Cryo-EM analysis of AaEnc

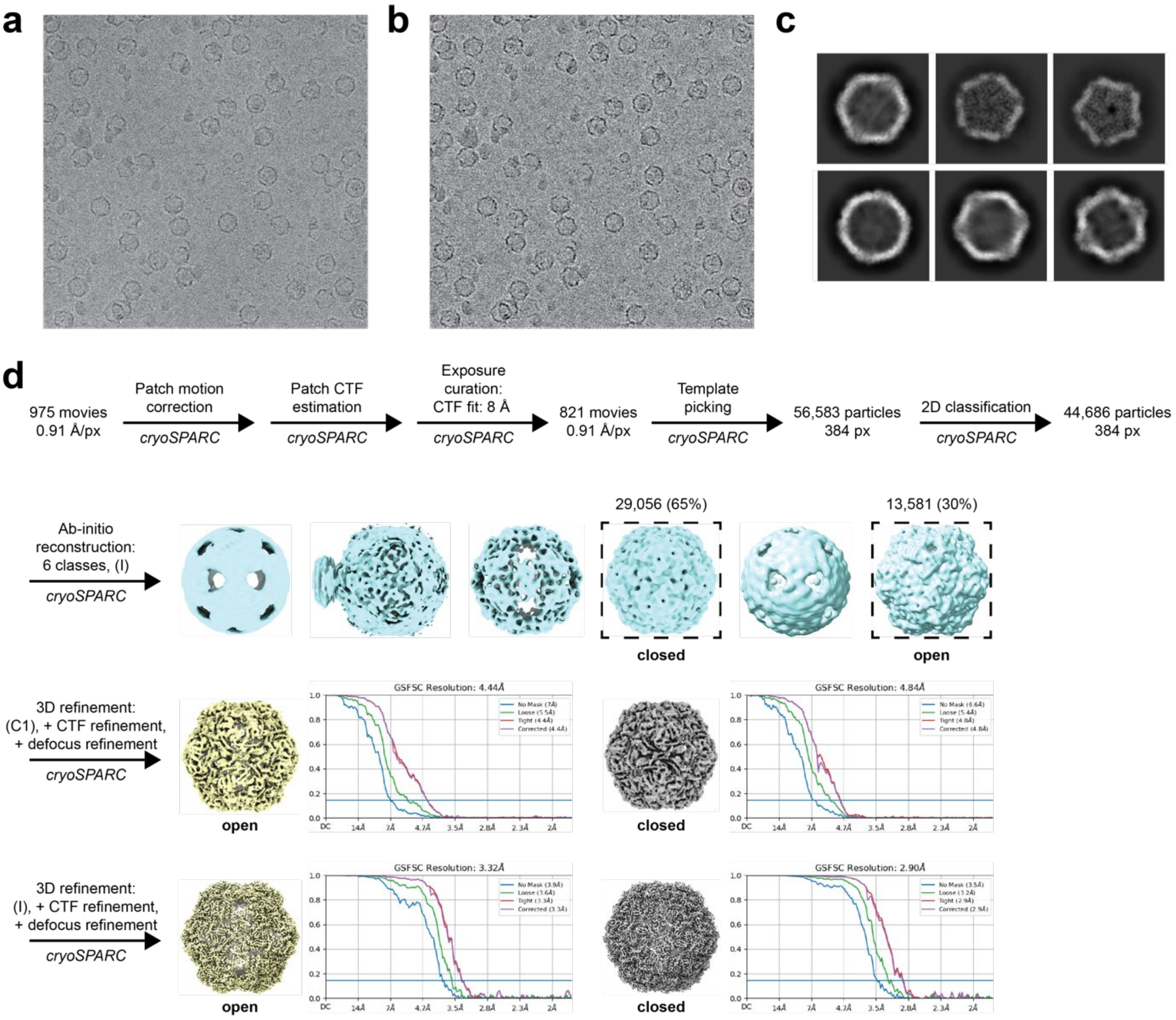

**Figure S3.** Cryo-EM analysis of AaEnc at pH 7.5.[9] a) Raw micrograph. b) Motion-corrected micrograph. c) Sample 2D class averages. d) cryo-EM data processing workflow.

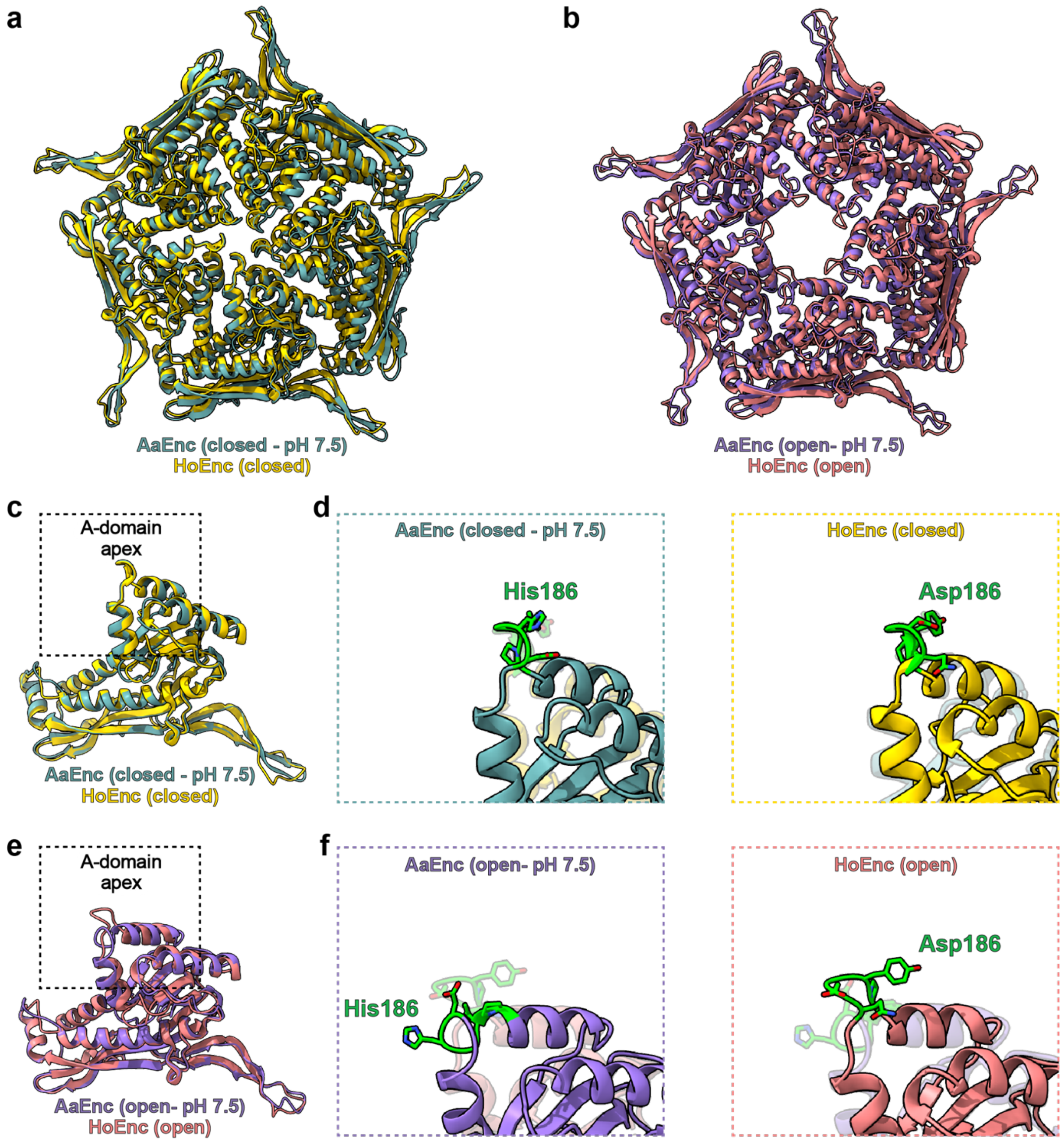

**Figure S4.** Structural comparison of the AaEnc at pH 7.5 and *H. ochraceum* 5-fold pores and pore loops. **a**) Top-down ribbon representation of the "closed" AaEnc pentamer (left, cyan) as compared to aligned ribbon representation of the "closed" *H. ochraceum* encapsulin pentamer (HoEnc; PDB 7OE2; left, yellow).[5] **b**) Top-down ribbon representation of the "open" AaEnc pentamer (right, purple) as compared to aligned ribbon representation of the "open" HoEnc encapsulin pentamer (PDB 7OEU; right, pink). **c**) Aligned and overlayed ribbon representation of the "closed" AaEnc protomer (cyan) and the "closed" HoEnc protomer (yellow; PDB 7OE2) with dashed box highlighting the A-domain. **d**) Magnified ribbon representation juxtaposing the A-domain of the "closed" AaEnc pentamer (cyan; left, solid; right, transparent) and the "closed" HoEnc protomer (yellow; right, solid; left, transparent), with corresponding loop residues of interest, including His186 for AaEnc and Asp186 for HoEnc (labeled), highlighted (green). **e**) Aligned and overlayed ribbon representation of the "open" AaEnc protomer (purple) and the "open" HoEnc protomer (pink; PDB 7OEU) with dashed box highlighting the A-domain. **f**) Magnified ribbon representation juxtaposing the A-domain of the "open" AaEnc pentamer (purple; left, solid; right, transparent) and the "open" HoEnc protomer (pink; right, solid; left, transparent), with corresponding loop residues of interest, including His186 for AaEnc and Asp186 for HoEnc (labeled), highlighted (green).

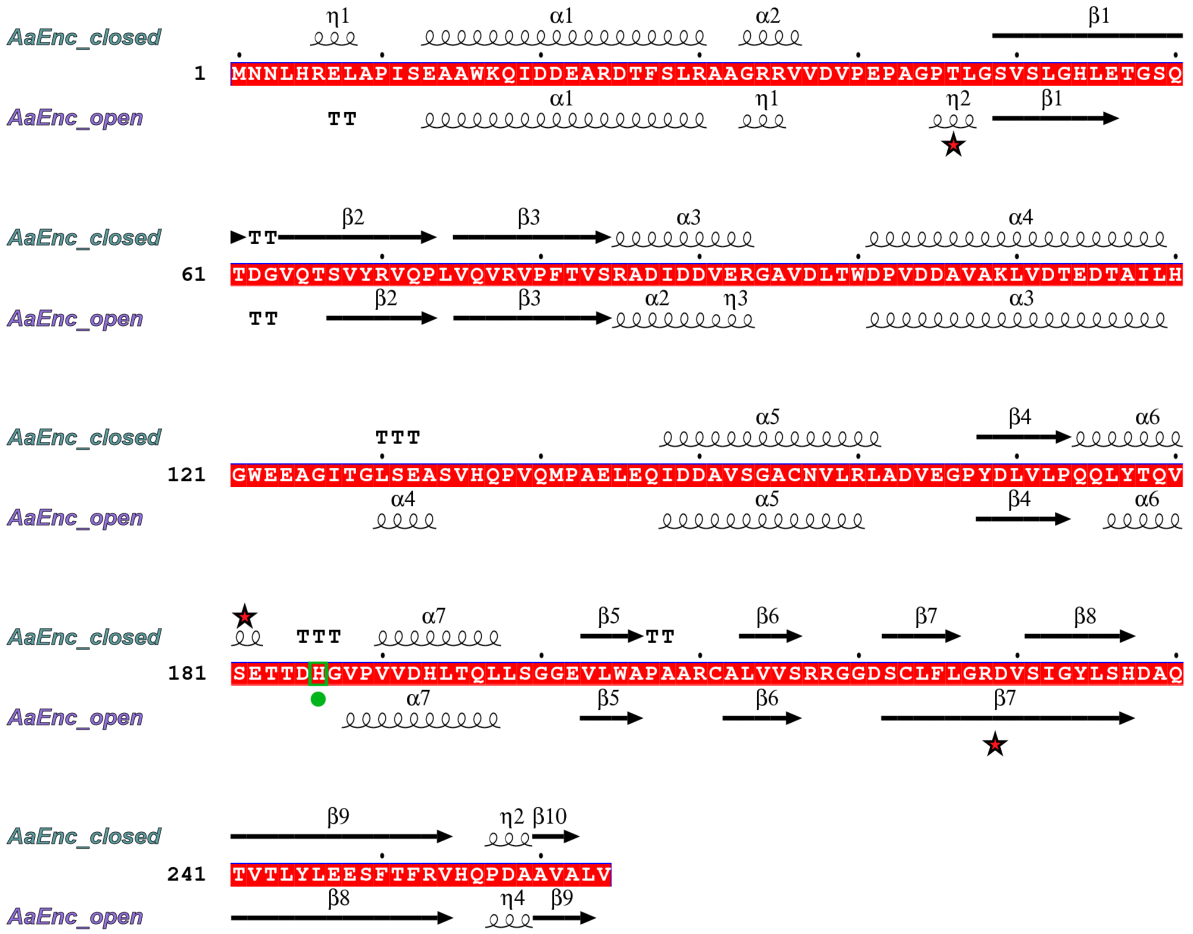

**Figure S5**. Sequence and secondary structure of AaEnc in the "closed" (top; cyan) vs "open" (bottom; purple) states at pH 7.5 generated with the ESPript 3 server (http://espript.ibcp.fr/).[7] Key differences are flagged, including the His186 residue highlighted with a green box and marked by a green circle. Additional discernible differences in secondary structure are marked with a red star. TT: β-turn; TTT: α-turn; α: α-helix; π-helix; β: β-strand.

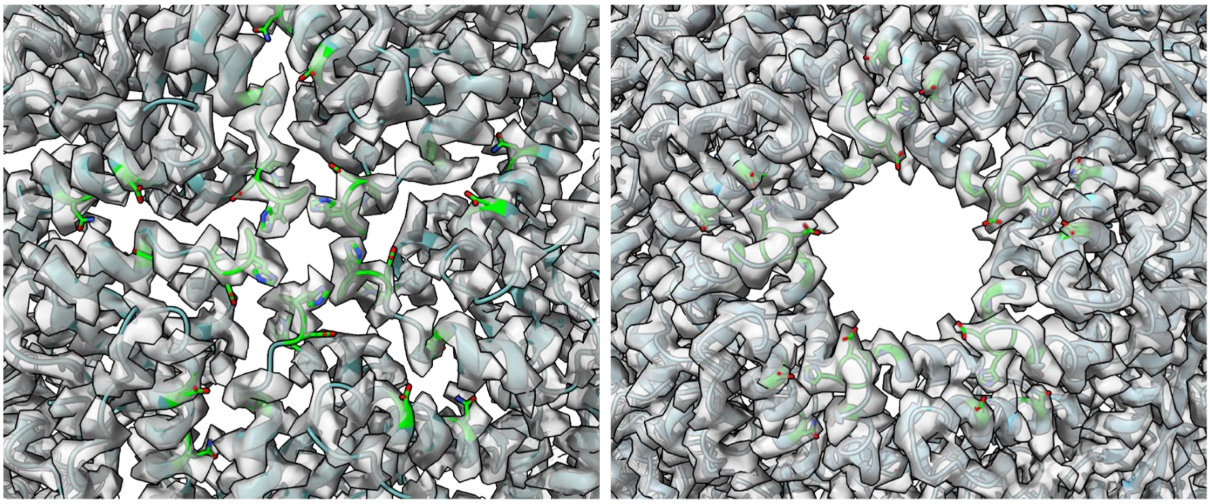

**Figure S6**. Cryo-EM density map to ribbon model comparison of AaEnc along the five-fold axis in the “closed” (left) and “open” (right) states at pH 7.5. Residues of interest, including Asp150, Asn157, Ser181, Asp185, His186, Gly187, Val188, and Pro189, highlighted in green with additional heteroatom stick representation. Models visualized using ChimeraX.[10]

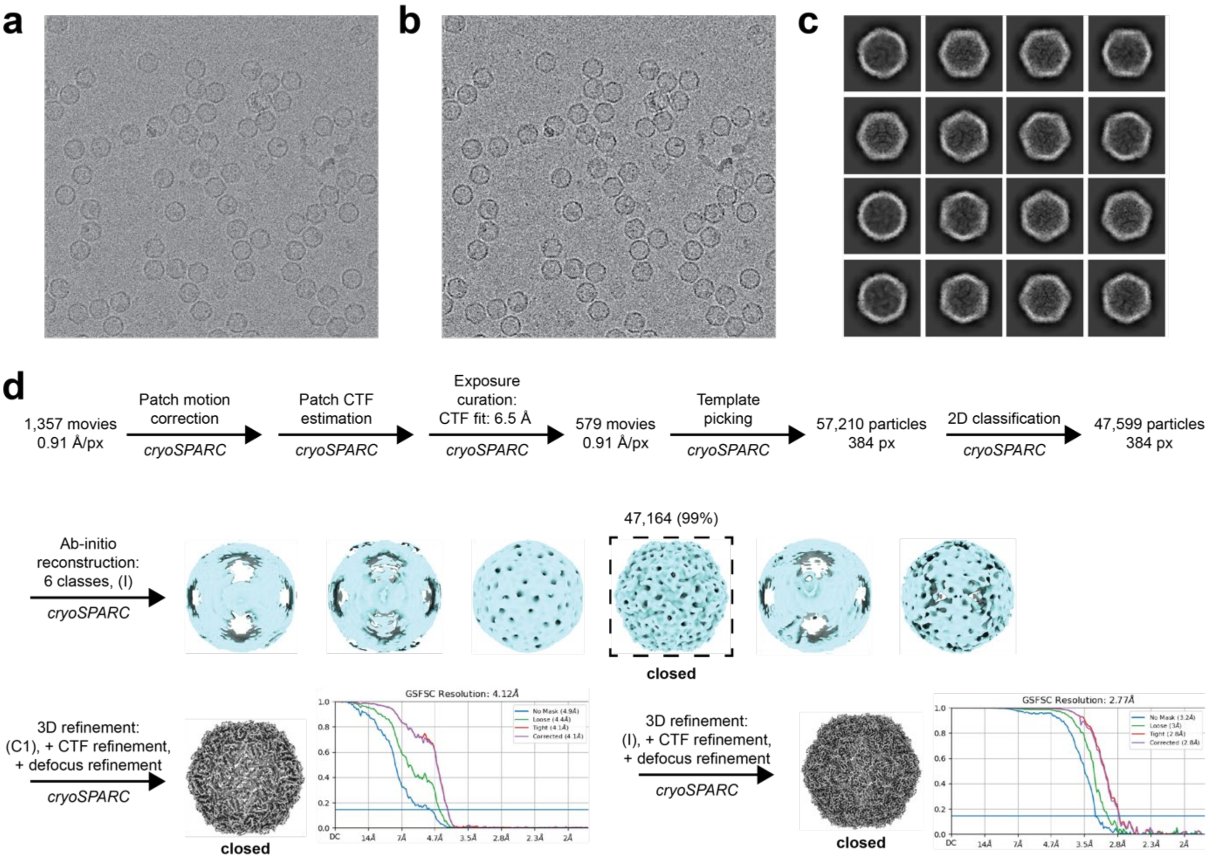

**Figure S7.** Cryo-EM analysis of AaEnc at pH 3.0.[9] a) Raw micrograph. b) Motion-corrected micrograph. c) Sample 2D class averages. d) Cryo-EM data processing workflow.

**Table S3**. Cryo-EM data collection and model building statistics.[9,11-15]

| Data Collection and Processing | pH 7.5 Closed | pH 7.5 Open | pH 3.0 Closed |
|---|---|---|---|
| **EMDB Accession** | EMD-27558 | EMD-27573 | EMD-27560 |
| **PDB Accession** | 8DN9 | 8DNL | 8DNA |
| Electron Microscope | Arctica | Arctica | Arctica |
| Voltage (kV) | 200 | 200 | 200 |
| Total dose ($e^-/Å^2$) | 41 | 41 | 43 |
| Defocus range (um) | -0.8 to -1.8 | -0.8 to -1.8 | -1.0 to -1.8 |
| Particles | 29,056 | 13,581 | 47,164 |
| Resolution (Å) | 2.90 | 3.32 | 2.77 |
| Map sharpening B-factor ($Å^2$) | -106.0 | -111.6 | -105.5 |
| Symmetry for map | I | I | I |
| Number of movies total | 975 | 975 | 1,357 |
| **Structure Refinement (ASU)** | | | |
| Structure B factor (mean) | 45.66 | 77.81 | 28.55 |
| Mean CC for protein (mask) | 0.83 | 0.79 | 0.83 |
| Chains in ASU | 1 | 1 | 1 |
| Bond lengths (Å) | 0.004 | 0.009 | 0.004 |
| Bond angles (°) | 0.993 | 1.259 | 0.941 |
| **Ramachandran Statistics** | | | |
| Favored % | 95.80 | 95.02 | 97.33 |
| Allowed % | 4.20 | 4.98 | 2.67 |
| Disallowed % | 0.00 | 0.00 | 0.00 |
| Rotamer outliers (%) | 0.45 | 0.92 | 0.00 |
| **MolProbity Score** | 1.54 | 1.90 | 1.38 |
| **Clashscore** | 4.78 | 10.92 | 4.80 |

### 6. AaEnc protomer structural alignments

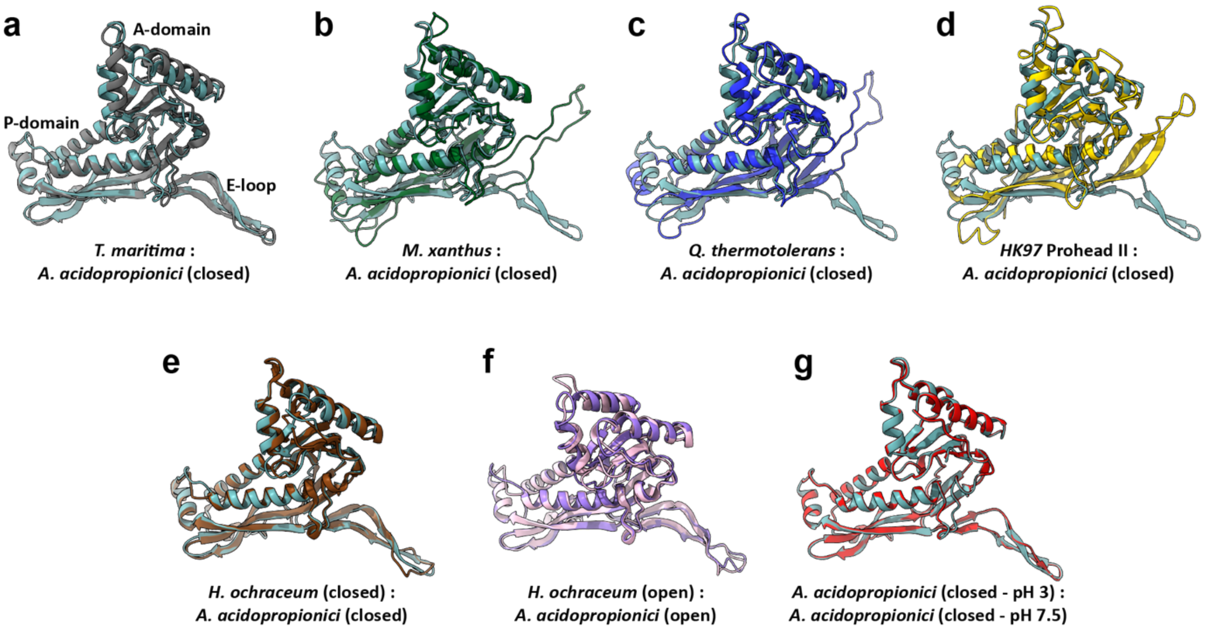

**Figure S8**. Structural alignments of the AaEnc protomer with other encapsulin protomers. A) AaEnc ("closed") alignment to the *Thermotoga maritima* encapsulin (PDB 3DKT). b) AaEnc ("closed") alignment to the *Myxococcus xanthus* encapsulin (PDB 4PT2). c) AaEnc ("closed") alignment to the *Quasibacillus thermotolerans* encapsulin (PDB 6NJ8). d) AaEnc ("closed") alignment to the HK97 Prohead II capsid protomer (PDB 3E8K). e) AaEnc ("closed") alignment to the *Haliangium ochraceum* closed encapsulin protomer (PDB 7OE2). f) AaEnc ("open") alignment to the *Haliangium ochraceum* "open" encapsulin protomer (PDB 7OEU).g) AaEnc ("closed") at pH 7.5 alignment to the AaEnc ("open") protomer at pH 3.0. Models visualized and alignments created using ChimeraX.[10]
